# Visual intracortical and transthalamic pathways carry distinct information to cortical areas

**DOI:** 10.1101/2020.07.06.189902

**Authors:** Antonin Blot, Morgane M Roth, Ioana T Gasler, Mitra Javadzadeh, Fabia Imhof, Sonja B Hofer

**Author notes:** Authors contributed equally (Antonin Blot, Morgane M Roth). Correspondence to: S.B.H.

## Abstract

Sensory processing involves information flow between neocortical areas, assumed to rely on direct intracortical projections. However, cortical areas may also communicate indirectly via higher-order nuclei in the thalamus, such as the pulvinar or lateral posterior nucleus (LP) in the visual system. The fine-scale organization and function of these cortico-thalamo-cortical pathways remains unclear. We find that responses of mouse LP neurons projecting to higher visual areas likely derive from feedforward input from primary visual cortex (V1) combined with information from many cortical and subcortical areas, including superior colliculus. Signals from LP projections to different higher visual areas are tuned to specific features of visual stimuli and their locomotor context, distinct from the signals carried by direct intracortical projections from V1. Thus, visual transthalamic pathways are functionally specific to their cortical target, different from feedforward cortical pathways and combine information from multiple brain regions, linking sensory signals with behavioral context.

## Introduction

Our perception of the environment is thought to rely on neuronal interactions within the cerebral cortex where sensory information is processed by hierarchical pathways involving many cortical areas (Van Essen, 1979). However, all cortical areas are also highly interconnected with the thalamus from which the cortex receives the majority of its input. First-order thalamic nuclei convey information from the sense organs to primary sensory areas in the neocortex and have been extensively characterized (Jones, 1985; Guillery and Sherman, 2002). However, the larger part of the sensory thalamus consists of so-called higher-order nuclei which form extensive and intricate circuits with cortical areas (Jones, 1985; Guillery and Sherman, 2002; Sherman, 2016). The complexity of these higher-order thalamo-cortical pathways makes it difficult to decipher their function.

The higher-order thalamic nucleus of the visual system is the pulvinar complex, known also as the lateral posterior nucleus (LP) in rodents (Roth et al., 2016; Baldwin et al., 2017; Zhou et al., 2017; Bennett et al., 2019). Pulvinar projections to primary visual cortex (V1) target mostly cortical layers 1 and 5a, and have been shown to convey contextual information (Roth et al., 2016) that can sharpen visual representations (Hu et al., 2019; Fang et al., 2020). However, the pulvinar provides more pronounced input to higher visual areas, where it also targets the cortical input layer 4 and can strongly impact cortical activity (Soares et al., 2004; Zhou et al., 2016; Beltramo and Scanziani, 2019). The pulvinar receives most of its input from visual brain areas. Some of its subdivisions are innervated by the superior colliculus, however, the main input to large parts of pulvinar comes from visual areas in the neocortex (Shipp, 2003; Roth et al., 2016; Baldwin et al., 2017; Zhou et al., 2017; Beltramo and Scanziani, 2019; Bennett et al., 2019). Therefore, this higher-order thalamic complex has been proposed to form transthalamic pathways, whereby layer 5 cortical cells of a lower-order area drive thalamocortical cells that project to a higher-order cortical area (Sherman and Guillery, 2011; Sherman, 2016). These indirect feedforward pathways via the thalamus would parallel direct intracortical feedforward connections, for instance from V1 to a higher visual area. While anatomical projection patterns are compatible with this hypothesis (Shipp, 2003; Bennett et al., 2019), the fine-scale input and output connectivity of pulvinar neurons has not been determined. It is therefore still unresolved if they are part of transthalamic, feedforward pathways between cortical areas. Alternatively, pulvinar circuits could provide additional visual pathways from the retina to the cortex via the superior colliculus or form specific, reciprocal loops with individual cortical areas (Wurtz et al., 2011; Guo Z. et al., 2017; Beltramo and Scanziani, 2019; Bennett et al., 2019; Guo K. et al., 2020). Furthermore, it is unclear what information these pathways through the pulvinar bring to cortical visual areas, and how the signals they convey differ from those carried by direct intracortical projections.

To address these questions, we focused on higher-order thalamic circuits of the mouse visual system. More than a dozen higher visual areas have been described in the mouse neocortex (Wang and Burkhalter, 2007; Zhuang et al., 2017), including the anterolateral area (AL) and the posteromedial (PM) area. The visual response properties of AL and PM are different from V1 and distinct from each other. The function of these visual areas is still unclear, but AL may be specialized to process visual motion, as neurons in AL preferentially respond to moving stimuli of low spatial and high temporal frequency, while PM neurons on average prefer high spatial and low temporal frequency stimuli (Andermann et al., 2011; Marshel et al., 2011; Roth et al., 2012; de Vries et al., 2020). Both areas receive prominent input from V1 and the mouse homologue of the pulvinar, LP (Wang and Burkhalter, 2007; Glickfeld et al., 2013; Roth et al., 2016; Bennett et al., 2019).

Using mono-synaptic rabies tracing, we found that the population of LP neurons projecting to either of these cortical areas combines information from V1 layer 5 cells with signals from many other cortical and subcortical areas, including superior colliculus. Two-photon calcium imaging of axonal boutons revealed that LP sends specific signals to higher visual areas that differ from those carried by the direct cortical feedforward pathway from V1. In behaving animals, direct projections from V1 to cortical area AL mainly carry information about visual motion in the environment, while LP input to AL combines information about visual motion and the animals’ own movement. Neurons imaged in AL showed very similar visual response properties to their input from LP, however, optogenetic silencing suggested that visual response properties of LP neurons targeting area AL were not inherited from AL. Our results therefore indicate that LP does not form exclusive reciprocal loops with cortical areas but is a key node of feedforward transthalamic pathways that convey specific visuo-motor information to higher visual areas, a contribution distinct from V1 intracortical feedforward projections. Thalamic pathways integrate visual input from V1 with information from many other brain areas, and may thus link sensory signals with the behavioral context in which they are encountered.

## Results

### LP neurons projecting to higher visual areas receive inputs from many cortical and subcortical areas

LP is interconnected with all visual areas. However, the sources of inputs to LP neurons projecting to a specific higher visual area are unknown. LP could potentially form strong reciprocal loops with the cortex, whereby thalamic neurons receive most of their input from their cortical target area (Guo Z. et al., 2017; Guo K. et al., 2020). Alternatively, LP could be part of transthalamic pathways whereby thalamic neurons projecting to a higher visual area receive their inputs from other structures, such as V1 or the superior colliculus (Sherman and Guillery, 2011; Sherman, 2016; Beltramo and Scanziani, 2019). To address this question, we performed projection-specific mono-synaptic rabies tracing (Figure 1, Figure S1) (Wickersham et al., 2007; Wall et al., 2010; Schwarz et al., 2015; Reardon et al., 2016). We targeted LP neurons projecting to higher visual area AL by expressing Cre-dependent TVA receptor and glycoprotein (G) in LP while retrogradely delivering Cre-recombinase via an injection of retroAAV-Cre in AL localized by intrinsic signal imaging (Figure S1A-C). Subsequently, glycoprotein-deleted rabies virus (RVdG) was injected into LP where it specifically infected AL-projecting neurons and spread presynaptically. Importantly, in the absence of Cre, no G or TVA expression was observed, ensuring that RVdG could not infect or spread to neurons non-specifically (Figure S2A-D).

**Figure 1:**
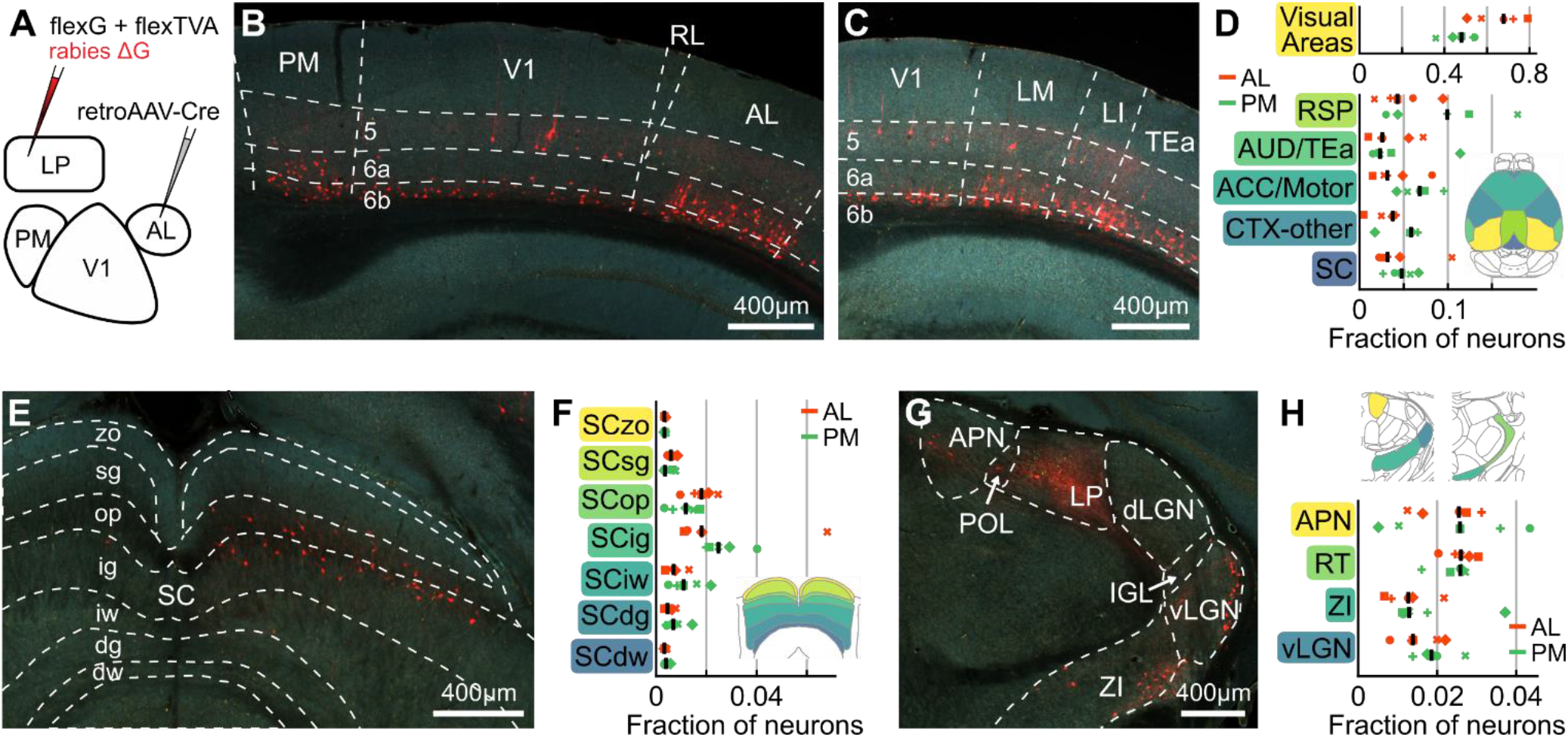
LP neurons projecting to higher visual areas receive input from many cortical and subcortical regions. **(A)** Schematic of the experimental design. RetroAAV-Cre was injected in cortical area AL or PM, then AAV-flex-G and AAV-flex-TVA were injected in LP. Finally, G-deleted pseudotyped N2C rabies virus was injected in LP, specifically labelling cells presynaptic to AL- or PM-projecting LP neurons. **(B,C)** Example images showing rabies-labelled neurons (red) presynaptic of AL-projecting LP neurons in visual areas of a coronal slice from a brain injected with retroAAV in AL. Numbers indicate the cortical layers. AL: anterolateral area, LI: latero-intermediate area, LM: lateromedial area, PM: posteromedial area, RL: rostrolateral area, TEa: temporal association areas, V1: Primary visual cortex. **(D)** Relative distribution of cells presynaptic to AL-projecting LP neurons (orange, retroAAV injection in AL, 5 mice) and PM-projecting LP neurons (green, retroAAV injection in PM, 5 mice) in fraction of total cells per brain. Symbols denote individual brains (similar across all plots). Black lines indicate median values. Inset: dorsal view of the mouse brain. The color-coding of highlighted areas matches that of their names along the y-axis. ACC/Motor: Anterior cingulate cortex and motor areas, AUD/TEa: auditory and temporal association areas, CTX-other: remaining cortical areas, SC: superior colliculus, RSP: retrosplenial cortex. **(E)** Example image of cells presynaptic to AL-projecting LP neurons in the superior colliculus of a brain with retroAAV injection in AL. SC: Superior colliculus, zo: zonal layer, sg: superficial gray layer, op: optic layer, ig: intermediate gray layer, iw: intermediate white layer, dg: deep gray layer, dw: deep white layer. **(F)** Distribution of cells presynaptic to AL-projecting LP neurons (orange) and PM-projecting LP neurons (green) across layers of the superior colliculus. Inset: coronal view of color-coded superior colliculus layers. See (E) for abbreviations. **(G)** Example image of cells presynaptic to AL-projecting LP neurons in inhibitory prethalamic and pretectal structures. APN: Anterior pretectal nucleus, dLGN: dorsal lateral geniculate nucleus, IGL: intrageniculate leaflet, LP: lateral posterior nucleus of the thalamus, POL: posterior limitans nucleus of the thalamus, vLGN: ventral lateral geniculate nucleus, ZI: zona incerta. **(H)** Distribution of presynaptic cells across inhibitory structures in the prethalamus and pretectum. Top shows coronal view of color-coded areas. APN: Anterior pretectal nucleus, RT: reticular thalamic nucleus, vLGN: ventral lateral geniculate nucleus, ZI: zona incerta.

We found that LP neurons projecting to AL received synaptic input from a large number of brain areas (Figure 1B-H, Figure S1E,F,H,I), resembling the general pattern of inputs to LP (Figure S2E-I; Roth et al., 2016; Bennett et al., 2019). Cells providing input to AL-projecting LP neurons were particularly abundant in visual cortical areas (Figure 1B-D, Figure 2A,B). Presynaptic neurons were also located in the ipsilateral superior colliculus (Figure 1E,F) as well as in cortical association areas, in particular the retrosplenial cortex and anterior cingulate and secondary motor cortices (Figure 1D, Figure S1E,F,H). Notably, LP neurons received input from several areas containing mainly inhibitory neurons, including the thalamic reticular nucleus, the zona incerta, the ventral lateral geniculate nucleus and anterior pretectal nucleus (Figure 1G,H) (Halassa and Acsády, 2016; Sabbagh et al., 2020), revealing LP as a target of multiple long-range inhibitory circuits. Therefore, populations of neurons in LP projecting to AL integrate signals from a variety of cortical and subcortical areas. To determine if this brain-wide pattern of input connectivity is specific to AL-projecting LP neurons or a general feature of LP thalamocortical pathways, we investigated the connectivity of LP neurons projecting to a different visual cortical area, PM. First, to ensure that LP projections to AL and PM originate from distinct populations of neurons, we injected differently colored retrograde tracers into the two cortical areas (Figure S3). In agreement with a previous study (Juavinett et al., 2020), we found that only a small subset of LP neurons was double-labelled (7.8 %; Figure S3F), indicating that the projections from LP to AL and PM are largely distinct. We then determined the sources of presynaptic input to PM-projecting LP neurons, employing projecting-specific mono-synaptic rabies tracing as described above, but with injection of retroAAV-Cre into PM (Figure S1G). PM-projecting LP neurons had a distribution of presynaptic inputs that was largely similar to that of AL-projecting neurons (Figure 1D,F,H, Figure S1H,I), suggesting that the pattern of inputs to LP thalamocortical pathways generalize across higher visual target areas. Notably, PM- and AL-projecting neurons had similar numbers of presynaptic cells in AL and PM (Figure 2A,B), suggesting that LP neurons do not preferentially receive reciprocal input from their cortical target area. Slight differences in the distribution of inputs to PM- and AL-projecting neurons were, however, apparent: PM-projecting neurons tended to be innervated to a larger extent by non-visual cortical areas (Figure 1D, Figure S1H) and deeper layers of SC (Figure 1F).

**Figure 2:**
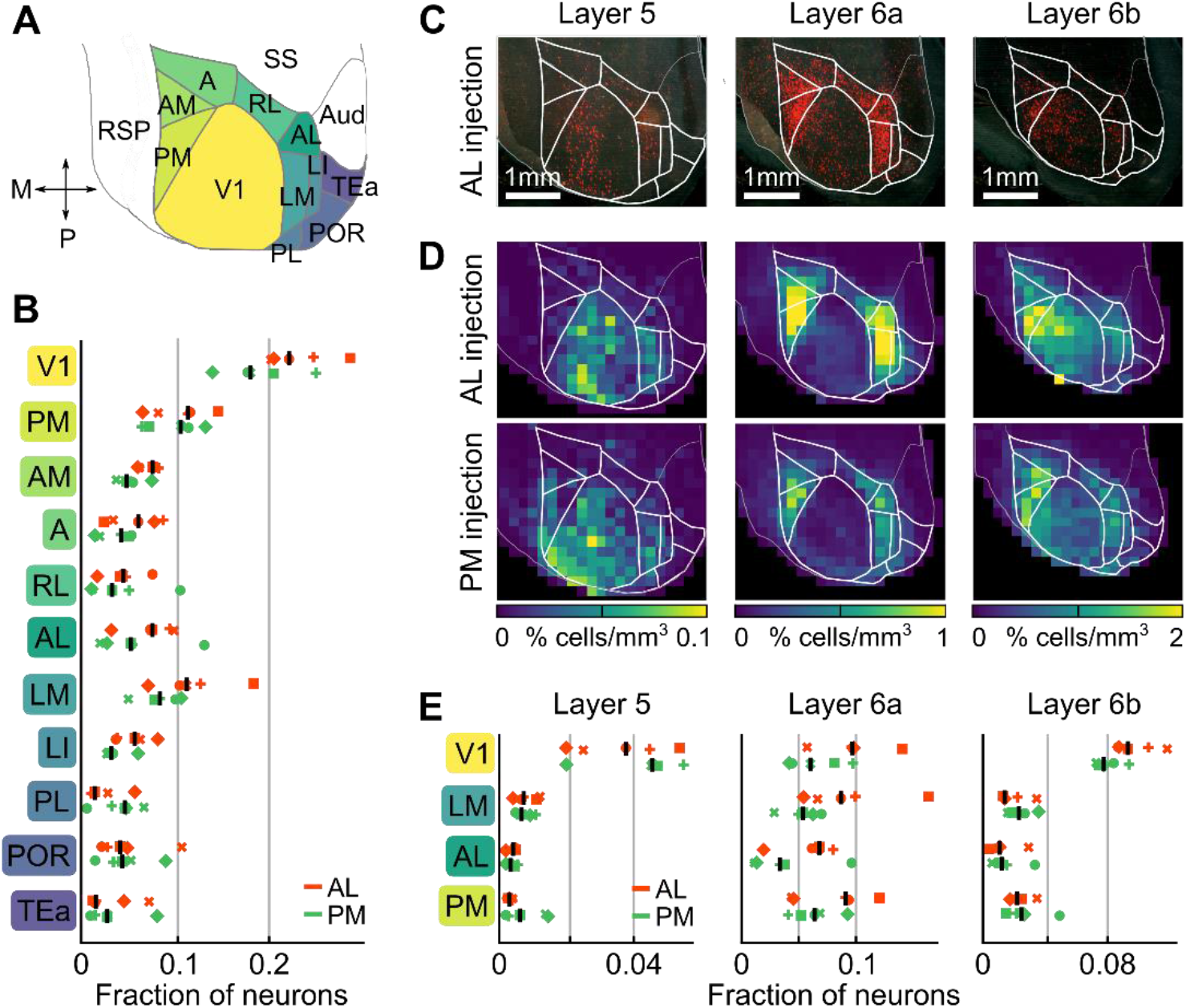
Distribution of cortical input to LP shows the hallmarks of feedforward transthalamic pathways. **(A)** Schematic dorsal view of cortical visual areas as shown in panels (C) and (D). The color-coding of the areas matches that of their names along the y-axis in (B) and (E). A: anterior area, AL: anterior lateral area, AM: anterior medial area, Aud: auditory areas, LI: laterointermediate area, LM: lateromedial area, PL: prelimbic area, PM: posteromedial area, POR: postrhinal area, RL: rostrolateral area, RSP: retrosplenial cortex, SS: somatosensory areas, TEa: temporal association areas, V1: primary visual cortex. **(B)** Fraction of cells in all visual areas shown in (A) presynaptic to AL-projecting LP neurons (orange, retroAAV injection in AL, 5 mice) and PM-projecting LP neurons (green, retroAAV injection in PM, 5 mice). Symbols denote individual brains (similar across all plots). Black lines indicate median values. See (A) for abbreviations. **(C)** Dorsal view of presynaptic cells in layers 5, 6a and 6b of an example brain with retroAAV injection in AL. The retroAAV injection site is visible in layer 5 (faint green DiO labelling). **(D)** Average relative density of presynaptic cells per volume in layer 5 (left), layer 6a (middle) and layer 6b (right) for all brains with retroAAV injection in AL (top) and PM (bottom). **(E)** Fraction of presynaptic cells in four cortical visual areas divided by layer. See (A) for abbreviations.

### Distribution of cortical input to LP shows the hallmarks of feedforward transthalamic pathways

Cortical efferents have been described to differentially affect their target neurons, depending on the cortical layer they originate from. The main driving input onto thalamic neurons from the cortex is thought to arise from layer 5 cells while layer 6 cells are assumed to provide weaker or modulatory feedback (Jones, 1985; Rockland, 1996; Crick and Koch, 1998; Sherman and Guillery, 2011; Sherman, 2016). To determine the cortical origin of putative driving and modulatory inputs onto AL- and PM-projecting LP neurons, we quantified the number of presynaptic cells in each layer of visual cortical areas (Figure 2C-E). We found that presynaptic layer 5 cells were not predominantly located in the cortical target area of either AL- or PM-projecting LP neurons but were by far most numerous in V1. In contrast, the density of presynaptic layer 6a cells was much higher in higher visual areas than in V1. Presynaptic layer 6b cells showed a distribution similar to layer 5 inputs to LP and may therefore represent a cell class distinct from layer 6a (Hoerder-Suabedissen et al., 2018). Together, these results suggest that reciprocal loops between higher visual areas and LP are not a dominant circuit motif of visual thalamocortical pathways. Instead, these pathways appear to have a strong feedforward component, whereby LP neurons integrate driving inputs from V1 layer 5 cells with information from many other cortical and subcortical areas.

### Thalamic and cortical inputs convey distinct visual information to higher visual areas

The above results indicate that LP neurons projecting to higher visual areas receive prominent feedforward input from V1. These feedforward transthalamic pathways parallel the direct feedforward intracortical projections from V1 to higher visual areas. However, it is unknown if intracortical and transthalamic pathways convey similar information to a cortical target area, or whether they are functionally distinct. To address this question, we used *in vivo* two-photon microscopy in awake, head fixed mice (Figure 3A), and imaged calcium signals of axonal projections from either LP or V1 within higher visual areas (Figure 3B-D, Figure S4A,B). We extracted fluorescence signals from micrometer-sized regions in cortical layer 1, corresponding to putative axonal boutons, and inferred spiking probability from calcium transients (Figure 3B,C; see Methods) (Petreanu et al., 2012; Glickfeld et al., 2013; Roth et al., 2016).

**Figure 3.**
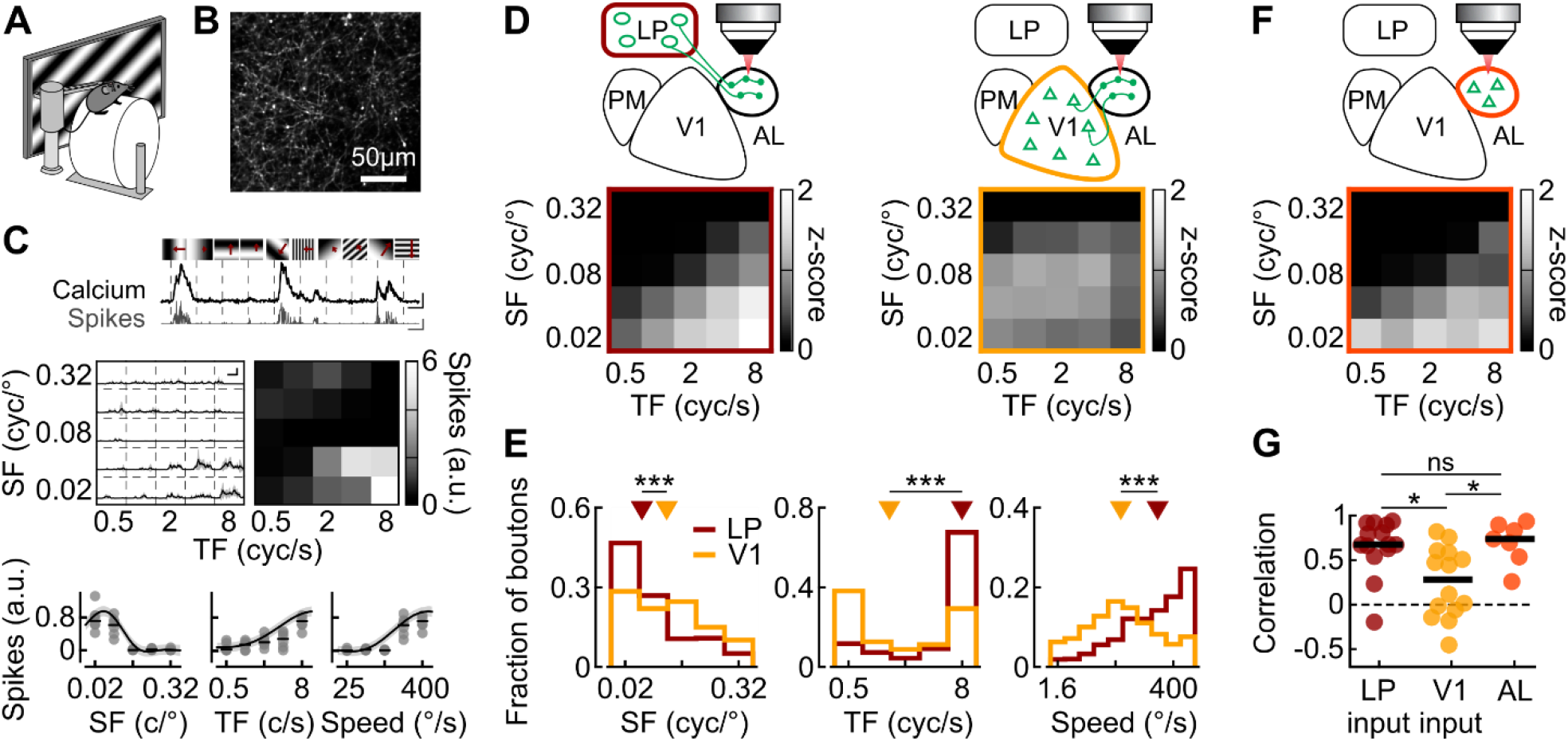
Thalamic and cortical inputs convey distinct visual information to higher visual area AL. **(A)** Schematic of the experimental design. Awake, head-fixed mice were presented with drifting gratings. **(B)** Example image of GCaMP6f expressing LP axons in cortical area AL. **(C)** Top: example ΔF/F traces (black, scale bars: 200 %, 2 s) and deconvolved traces of extracted spike rate (grey, scale bars: 2.5 a.u., 2 s). Gratings of various orientations and spatial frequencies (indicated on top) were presented and kept static for 1.2 s before drifting (onset indicated by the dotted lines) at various temporal frequencies (indicated by the length of the red arrows) for 2.2 s. Middle left: mean inferred spike rate across trials for the same bouton shown on top in response to gratings of different temporal frequencies (TF) and spatial frequencies (SF). Grey shading: standard deviation. Scale bar 1 s and 1 a.u. Middle right: response matrix obtained by averaging responses during the time of grating drift. Bottom: spatial frequency (left), temporal frequency (middle) and speed (right, ratio between TF and SF) response curves of the same bouton. Grey dots represent single trials, black dashes depict medians, black curves and grey shading are predictions from the GP fit of the response and their standard deviation (see Methods). **(D)** Top: Schematic of the experimental design. GCaMP6f was expressed either in LP (left) or V1 (right) and axonal boutons were imaged in AL. Bottom: matrix of average population responses to gratings of different temporal frequency (TF, x-axis) and spatial frequency (SF, y-axis) of LP boutons (left) and V1 boutons (right) in AL. 3732 and 3371 boutons, from 14 and 14 sessions in 14 and 7 mice for LP and V1 boutons respectively. **(E)** Distribution of preferred spatial frequency (left, 2237 LP and 2555 V1 boutons modulated by spatial frequency) preferred temporal frequency (middle, 1333 LP and 1637 V1 boutons modulated by temporal frequency) and preferred speed (2468 LP and 2928 V1 boutons modulated by speed) of LP boutons (dark red) and V1 boutons (yellow) recorded in AL. Triangles indicate the median. All p-values < 10^−50^. **(F)** Same as (D) for neurons recorded in AL (265 neurons from 7 sessions in 5 mice). **(G)** Pearson correlation coefficient between the average response matrix of the population of AL neurons shown in (F) and the population response matrices of individual recording sessions of LP boutons in AL (dark red), V1 boutons in AL (yellow) and AL neurons (orange). Circles represent individual sessions. Black lines depict medians. *: P < 0.01; ***: P < 10^−50^.

We first determined the visual response properties of LP boutons recorded in AL by presenting drifting gratings with different spatial and temporal frequencies and of varying orientations (Figure 3C). To visualize the spatio-temporal tuning of LP input to AL at the population level, we averaged the z-scored responses of all single boutons to each combination of spatial and temporal grating frequency at their preferred grating direction, resulting in a spatial and temporal frequency population response matrix (Figure 3D, left). We then compared the population response of LP boutons to that of direct intracortical projections from V1, by measuring visual responses of V1 boutons in the same cortical area AL (Figure 3D, right). Population response matrices of LP and V1 boutons were markedly different. LP population activity was strongest in response to stimuli with low spatial and high temporal frequency, while V1 population activity was less specific, responding to a wider range of stimuli. To determine whether this difference was apparent at the single bouton level, we fitted bouton responses with a Gaussian process regression model (see Methods) (Kim et al., 2018). We thus obtained single bouton tuning curves for temporal and spatial frequencies as well as for the ratio between the two: the speed of the grating drift (Figure 3C, bottom). Both V1 and LP boutons in AL had diverse spatio-temporal frequency preferences. LP boutons in AL preferred stimuli with lower spatial frequency, higher temporal frequency and therefore higher speed than V1 boutons (Figure 3E, all p-values < 10^−50^, Wilcoxon rank-sum test, all p-values for data grouped by recording session < 0.01, see Figure S4F-H). LP thus conveys specific visual information to AL, different from the visual signals carried by V1 projections to the same cortical area.

To determine if the above results were specific for cortical area AL or if transthalamic and intracortical pathways are in general functionally distinct, we repeated our experimental protocol while imaging LP and V1 boutons in higher visual area PM. We found that the population response of LP boutons in PM was again different from that of V1 boutons recorded in the same area (Figure 4A). Individual LP boutons responded best to stimuli of higher temporal frequency, lower spatial frequency and higher speed than V1 boutons in the same area (Figure 4B, all p-values < 10^−30^, all p-values for data grouped by recording session < 0.04, see Figure S4F-H), similar to what we had observed for LP and V1 boutons in AL. Therefore, cortical and thalamic inputs convey distinct visual information to the same higher visual area.

**Figure 4.**
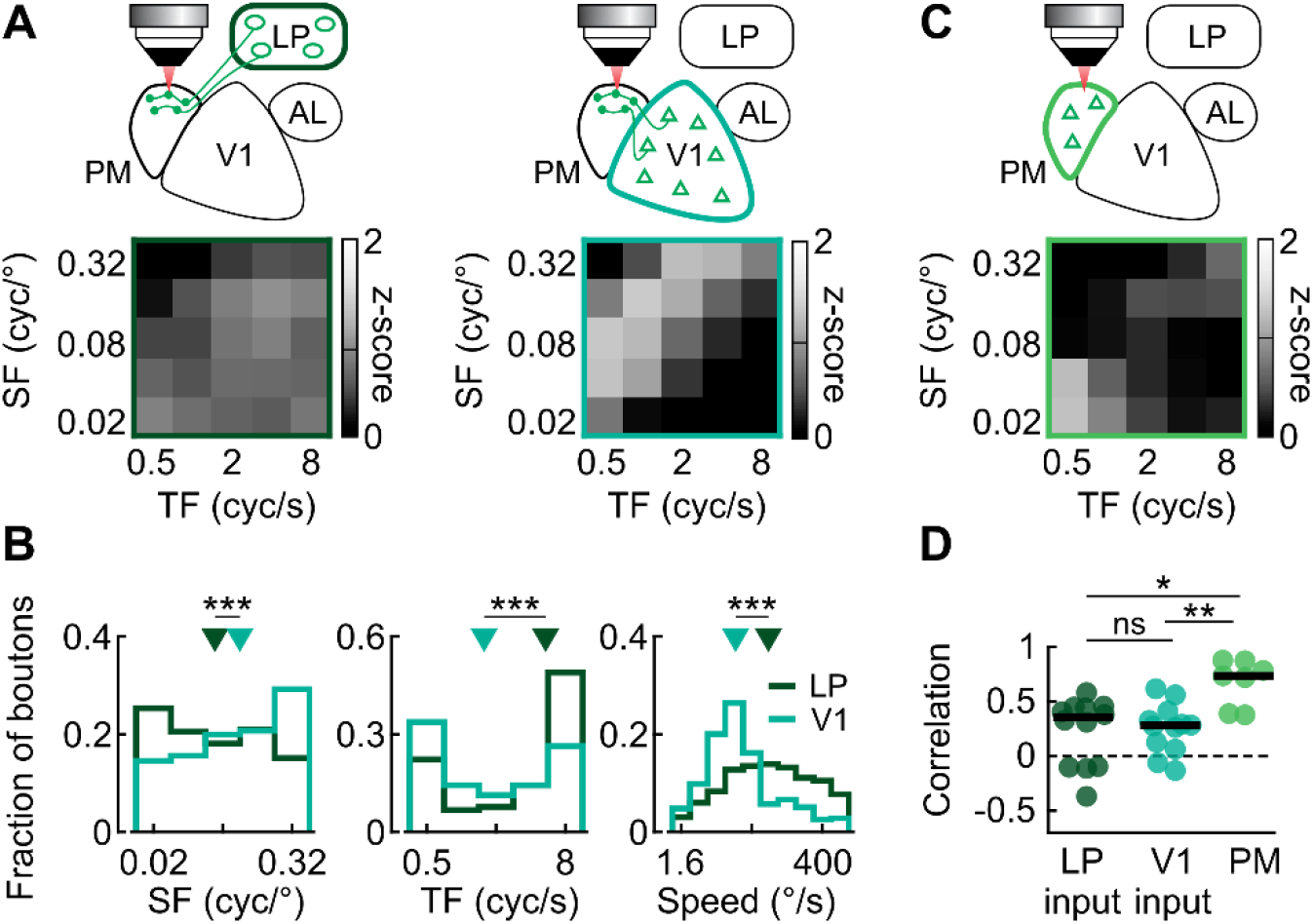
Activity in area PM shows aspects of both V1 and LP input. **(A)** Top: Schematic of the experimental design. GCaMP6f was expressed either in LP (left) or V1 (right) and axonal boutons were imaged in cortical area PM. Bottom: matrix of average population responses to gratings of different temporal frequency (TF, x-axis) and spatial frequency (SF, y-axis) of LP boutons (left) and V1 boutons (right) in PM. 3361 and 3235 boutons, from 12 and 12 sessions in 10 and 7 mice for LP and V1 respectively. **(B)** Distribution of preferred spatial frequency (left, 2059 LP and 2327 V1 boutons modulated by SF), preferred temporal frequency (1128 LP and 1659 V1 boutons modulated by TF) and preferred speed (2342 LP and 2535 V1 boutons modulated by speed) of LP boutons (dark green) and V1 boutons (blue) recorded in PM. Triangles indicate medians. All p-values <10^−30^. **(C)** Same as (A) for neurons recorded in PM (341 neurons from 8 sessions in 5 mice). **(D)** Pearson correlation coefficient between the average response matrix of the population of AL neurons shown in (C) and the population response matrices of individual recording sessions of LP boutons in PM (dark green), V1 boutons in PM (blue) and PM neurons (green). Circles represent individual recording sessions; black lines depict medians. *: P <0.01; **: P <0.005; ***: P <10^−30^.

LP projections to different visual areas consistently preferred higher temporal and lower spatial frequency stimuli than V1 projections. However, the information conveyed by LP projections was nevertheless specific to their cortical target area. As previously shown for V1 inputs into AL and PM (Glickfeld et al., 2013), LP boutons in AL and PM showed significantly different visual response properties, with LP boutons in AL preferring gratings of higher temporal frequency, higher speed and lower spatial frequency than LP boutons in PM (Figure S4C, all p-values < 10^−20^, all p-values for data grouped by recording session < 0.007, see Figure S4F-H). Moreover, tuning curves of LP boutons to different visual stimulus features were comparable to those of V1 boutons or neurons in the same area (Figure S4I-K). These results indicate that different higher visual areas receive distinct information from LP, tuned to specific visual features.

### AL neurons and thalamic inputs to AL share similar response properties

Our results show that transthalamic and intracortical pathways converging on the same visual area carry distinct visual information. This raises the question of how the response properties of these two pathways relate to those of their target areas. We therefore measured visual response properties of neurons in cortical areas AL and PM and compared them to the properties of LP and V1 inputs to these areas. Surprisingly, in area AL, the population response of cortical neurons to gratings of different spatial and temporal frequencies (Figure 3F) was more similar to LP than to V1 input (Figure 3D). We quantified this similarity as the correlation coefficient between the average population response matrix of all AL neurons and that of individual recording sessions of AL neurons, LP boutons or V1 boutons (Figure 3G). The correlation between the population response of LP boutons in different recording sessions and the average AL population response matrix was high (Figure 3G; 0.65 +/− 0.31), in fact as high as when comparing individual recording sessions of AL neurons with the AL population average (Figure 3G; 0.70 +/− 0.24; LP vs AL, p = 0.7). In contrast, the population responses of V1 boutons in AL were poorly correlated with the AL population response matrix (Figure 3G; 0.25 +/− 0.40; AL vs V1: p = 0.01, LP vs V1: p = 0.003). Analyzing response properties of individual boutons and neurons revealed that AL neurons and LP boutons in AL were particularly well matched in their spatial frequency preferences, while AL neurons showed distributions of temporal frequency and speed preferences that lay in between distributions of V1 and LP boutons in AL (Figure S4D,F-H). These results suggest that while the population response in AL is better matched to LP input, AL may integrate information from both LP and V1.

Such potential integration of thalamic and cortical inputs by their target area was more consistent with activity in PM. Population responses of PM neurons diverged from both LP and V1 input but encompassed aspects of both (Figure 4A,C,D). Individual PM neurons were better matched to V1 boutons in their temporal frequency preferences, but better matched to LP boutons in their spatial frequency preferences (Figure S4E-H). These results indicate that LP input consistently carries visual information about higher temporal frequencies while V1 input conveys information about higher spatial frequencies relative to their cortical target areas. These distinct signals from thalamic and intracortical projections might be combined in various ways by different higher visual areas in the cortex.

### LP neurons do not inherit their tuning from their target area

Our results show that LP boutons in AL have similar visual response properties to their target area AL (Figure 3). This could be a result of particularly strong reciprocal loops between these areas, such that response properties of LP neurons projecting to AL are inherited from AL. The results from mono-synaptic rabies tracing suggest otherwise (Figure 2), however, this technique only gives an indication of anatomical connectivity, not of the functional influence of presynaptic inputs. To test the influence of AL activity on the visual response properties of LP neurons projecting to AL, we optogenetically suppressed AL by activating a depolarizing opsin (the channelrhodopsin variant Chrimson) (Klapoetke et al., 2014) in parvalbumin-positive interneurons while simultaneously imaging visual responses of LP boutons in AL (Figure 5, Figure S5A). This manipulation abolished most activity across all layers in AL (Figure S5B,C).

**Figure 5.**
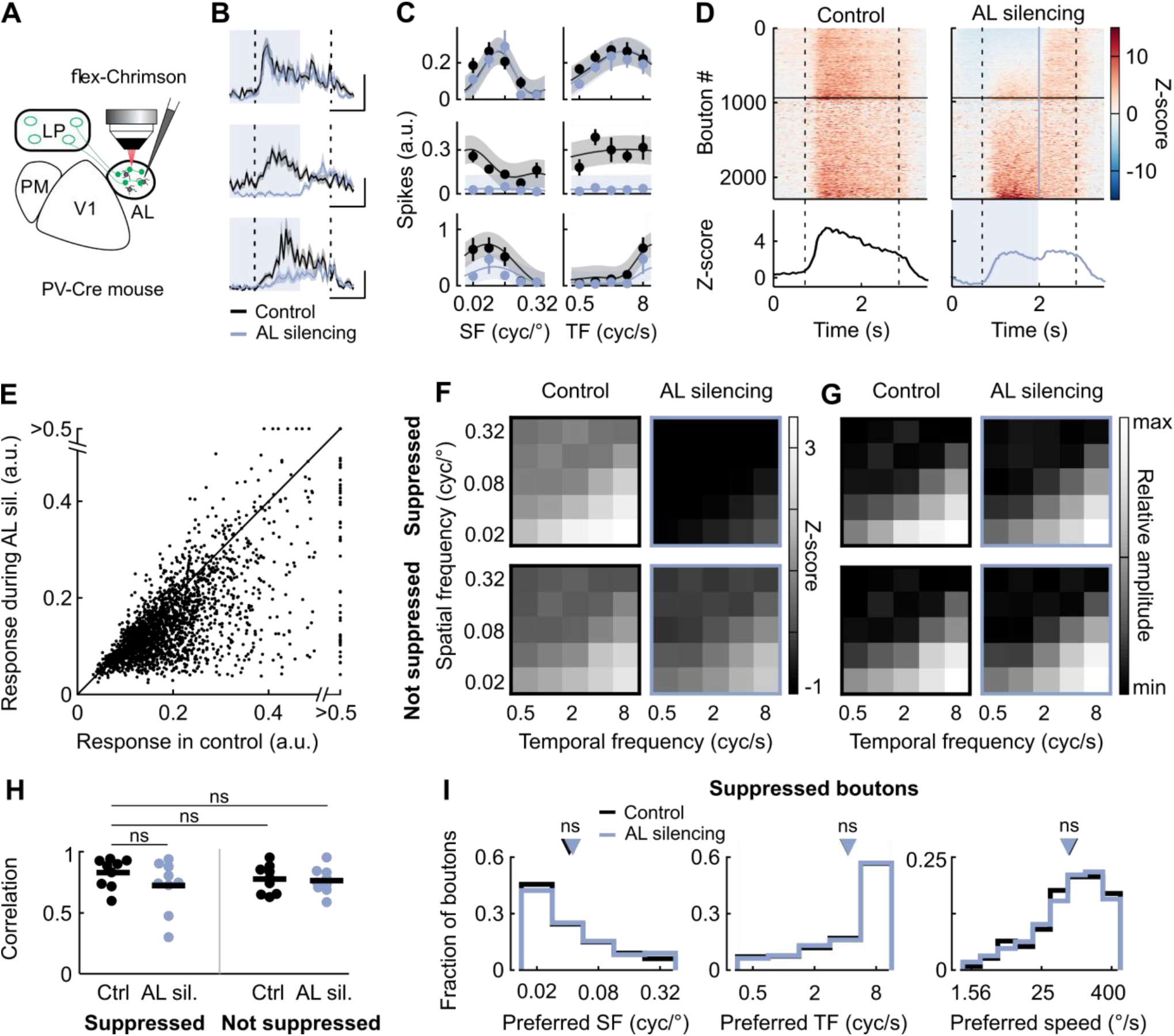
LP neurons projecting to AL do not inherit their visual response properties from AL. **(A)** Schematic of the experimental design. GCaMP6f was expressed in LP, while parvalbumin (PV) positive cells in AL expressed Chrimson. Two-photon imaging of LP boutons was performed in AL while a 637-nm laser illuminated AL to optogenetically activate PV cells and suppress local cortical activity. **(B)** Average responses of three example LP boutons in control trials (black) and during silencing of AL (blue) aligned to the onset of the drifting grating. Dotted vertical lines indicate the duration of grating presentation. Blue shading indicates the time of optogenetic activation. Grey shading indicates the standard error of the mean (sem). Scale bars: top, 1 s and 0.2 a.u.; middle, 1 s and 0.2 a.u.; bottom, 1 s and 0.5 a.u. **(C)** Spatial frequency (SF, left) and temporal frequency (TF, right) response curves for the boutons shown in (B). Black and blue dots indicate the average response in control and AL silencing trials, respectively. Error bars indicate sem. Curves and shading indicate predictions of the GP fit and standard deviation. **(D)** Top: time course of the z-scored activity of individual boutons. For each bouton, activity was averaged across grating stimuli evoking a response (see Methods) in control trials (left) and AL silencing trials (right). Responses are aligned to the onset of the laser. Boutons significantly suppressed by cortical silencing are shown at the top. Vertical lines indicate the duration of grating presentation. The blue line indicates the end of the optogenetic activation. Bottom: averaged z-scored activity across all boutons. Blue shading indicates the time of AL silencing. Grey shading indicates sem. **(E)** Relationship between the average moving grating responses of individual boutons with and without AL silencing (2299 boutons from 9 sessions in 6 mice). Sil: Silencing. **(F)** Average spatial and temporal frequency population response matrices as in Figure 3D for LP boutons that were suppressed (top) or not suppressed (bottom) by AL silencing, in control trials (left) and in silencing trials (right). **(G)** Same as (F) but in relative amplitude, normalized within each matrix. **(H)** Pearson correlation coefficient between the average response matrix of suppressed boutons in control trials (top left in F) and response matrices of individual sessions. Circles denote individual sessions. Black lines depict medians. Ctrl: control trials; AL sil: trials with AL silencing. **(I)** Distribution of preferred spatial frequency (left, 610 boutons), preferred temporal frequency (middle, 337 boutons) and preferred speed (right, 731 boutons) for boutons suppressed by AL silencing in control trials (black) and in AL silencing trials (blue). Triangles indicate the median. In all panels data from 9 sessions in 6 mice.

Silencing AL decreased visually-evoked activity in a large fraction of LP boutons in AL (39 +/− 15 % of boutons in each session, 55 +/− 22 % decrease in response amplitude of suppressed boutons; mean +/− standard deviation, Figure 5B-E). This decrease in activity was not observed in control animals without opsin expression (Figure S5D-F). Therefore, AL provides significant input onto LP neurons projecting back to this cortical area. However, the response properties of LP boutons in AL were not altered by AL silencing (Figure 5F-I, Figure S5G-J). Boutons whose responses were suppressed during AL silencing showed residual visual preferences and population responses similar to their responses without silencing (Figure 5F-I). In addition, not only the response preferences of LP boutons but also their response specificity were unaffected by AL silencing, since spatial and temporal frequency tuning widths were similar with and without AL silencing (Figure S5J). Furthermore, LP boutons suppressed by AL silencing were not different from the rest of the population. Indeed, in control trials without optogenetic stimulation, boutons suppressed by AL silencing and unaffected boutons showed similar responses to gratings of different spatial and temporal frequencies (Figure 5F-H, Figure S5H, all p-values > 0.1), although suppressed boutons were slightly more active (Figure S5H). Neither the fraction of affected neurons, nor the strength of suppression were correlated with the response preferences of LP boutons (Figure S5I). Therefore, while AL exerts a strong influence on LP activity, the visual response properties of LP neurons projecting to AL are not inherited from AL, but closely resemble responses of AL neurons even when the input from this cortical area is removed.

### Thalamic and cortical inputs convey distinct visuo-motor information to higher visual areas

Our results indicate that LP is a key node in transthalamic pathways which send specific visual information to higher visual cortical areas. Other studies additionally suggest that transthalamic pathways convey non-visual information (Komura et al., 2013; Saalmann and Kastner, 2015). We have previously shown that LP sends visual and non-visual contextual signals to V1 that carry information about visual scene changes not predicted by the animal's own actions (Roth et al., 2016). LP may therefore integrate visual with contextual information, for instance about the movement of the animal. To explore this possibility, we used two-photon calcium imaging of LP boutons in AL and compared their responses to V1 boutons imaged in the same cortical area in head-fixed mice running on a cylinder through a virtual environment (Figure 6A). We habituated mice to a virtual linear corridor where the motion of visual patterns displayed on monitors was controlled by the running speed of the animal, similar to previous studies (Poort et al., 2015; Roth et al., 2016). During calcium recordings we then uncoupled the virtual optic flow from the animals’ locomotion by replaying movies of the virtual corridor recorded in previous sessions (see Methods). This allowed us to separately assess the effect of locomotion and visual motion on neuronal activity (Figure 6B,C). We quantified how strongly neuronal activity inferred from calcium signals was modulated by either running speed or optic flow by computing mean cross-correlation coefficients between neuronal activity and these variables for each bouton (average over lags spanning 250 ms, see Methods). These correlation coefficients revealed diverse relationships, including positive and negative correlations (Figure 6B,C).

**Figure 6.**
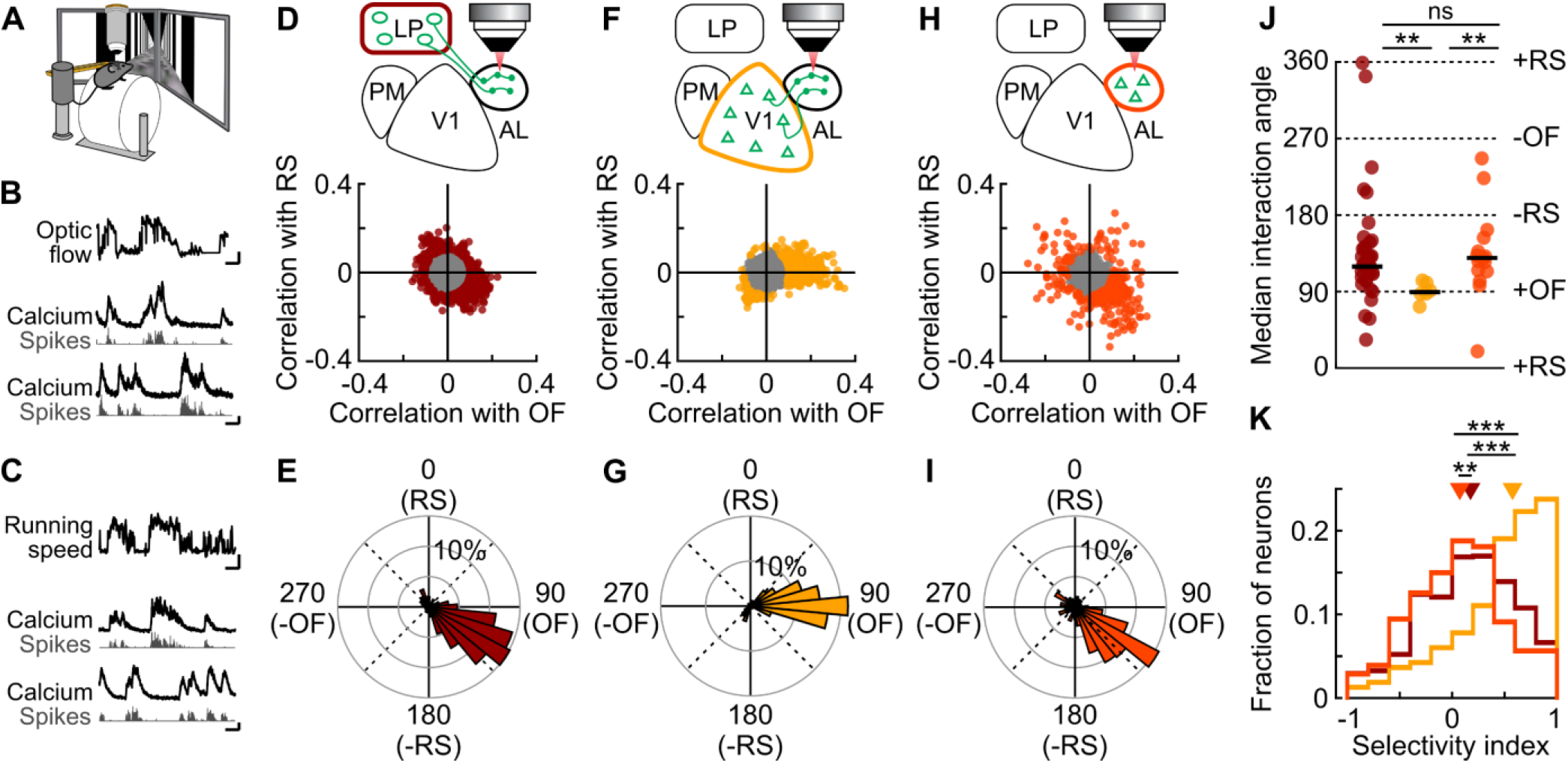
Thalamic and cortical inputs convey distinct visuo-motor information to higher visual areas. **(A)** Schematic of the experimental design. Two-photon imaging in a head-fixed mouse running through a virtual corridor allowing optic flow to be uncoupled from the animal’s locomotion while recording neuronal responses. **(B)** Top trace: optic flow speed; below: example activity traces of two AL neurons positively (middle, correlation coefficient R = 0.16) and negatively correlated (bottom, R = −0.23) with optic flow. Black, calcium trace; grey, inferred spikes. Scale bar for optic flow, 5 s and 10 cm/s, scale bar for neuronal activity, 5 s and 100% ΔF/F/ 1 a.u. **(C)** Top trace: running speed; below: example activity traces of two AL neurons positively (middle, R = 0.24) and negatively correlated (bottom, R = −0.23) with running speed. black, calcium trace; grey, inferred spikes. Scale bar for running speed, 5 s and 10 cm/s; scale bar for neuronal activity, 100% ΔF/F/ 1 a.u. **(D)** Top: schematic of the recording configuration. GCaMP6f was expressed in LP while calcium activity of LP boutons in AL was recorded using two-photon imaging. Bottom: relationship between the mean cross-correlation coefficients (see Methods) of neuronal activity with running speed and optic flow speed for all responsive LP boutons (12369 boutons from 43 sessions in 18 mice). Only boutons with mean cross-correlation greater than 0.1 (colored points in scatter plot) were included in the analysis shown in (E), (J) and (K). **(E)** Histogram in polar coordinates showing the distribution of interaction angles between the mean cross-correlation of activity with running speed (RS) and optic flow speed (OF) for LP boutons imaged in AL (919 boutons from 43 sessions in 18 mice). **(F)** Same as (D) for V1 boutons imaged in AL (2363 boutons from 6 sessions in 3 mice). **(G)** Same as (E) for V1 boutons imaged in AL (338 boutons from 6 sessions in 3 mice). **(H)** Same as (D) for cortical neurons imaged in AL (735 neurons from 15 sessions in 5 mice). **(I)** Same as (E) for cortical neurons imaged in AL (289 neurons from 15 sessions in 5 mice). **(J)** Median interaction angles across single sessions for LP boutons in AL (left), V1 boutons in AL (middle) and AL neurons (right). **: P < 0.01; ns: non-significant, P = 0.13. **(K)** Distribution of selectivity indices (difference of the absolute mean cross-correlation of neuronal activity with optic flow and running speed divided by their sum) for individual LP boutons in AL (dark red), AL neurons (orange) and V1 neurons in AL (yellow). −1 indicates a high correlation only with running speed, 1 a high correlation only with optic flow and 0 indicates equally high correlation with running speed and optic flow of individual boutons/neurons. **: P < 10^−3^; ***: P < 10^−27^.

To estimate the degree to which information about visual motion and running speed is integrated at the level of individual boutons, we plotted the mean correlation coefficients between neuronal activity and the two variables against each other for each bouton (Figure 6D,F). For boutons carrying information about running speed or optic flow (mean cross-correlation coefficient >= 0.1) we then derived an interaction angle θ (see Methods). Values of θ close to 0° indicate a bouton not modulated by optic flow speed, but positively correlated with running speed, increasing its responses with increasing running speed. Values close to 180° indicate a bouton whose activity was negatively correlated with running speed and decreased its responses with increasing running speed. Accordingly, θ values of 90° or 270° indicate that a bouton was not modulated by running speed, but its activity was positively or negatively correlated with optic flow speed, respectively. Finally, oblique angles correspond to neurons informative about both variables.

The large majority of V1 boutons recorded in cortical area AL had interaction angles around 90°, denoting that their activity was positively correlated with optic flow speed but not correlated with running speed (Figure 6G). Therefore, while the activity of V1 neurons has been shown to be modulated by locomotion (Niell and Stryker, 2010), under our experimental conditions, V1 projections to AL convey mainly visual information about the speed of visual motion. In contrast, the activity of many LP boutons was modulated both by optic flow and running speed. Neuronal activity in these boutons was predominantly positively correlated with optic flow speed and negatively correlated with running speed, indicating that they were activated by visual motion but suppressed by running (Figure 6D,E). Accordingly, LP boutons exhibited mainly interaction angles between 90° and 180°, significantly different from the distribution of angles of V1 boutons (Figure 6J). These data suggest that while V1 projections to AL mostly provide a channel for visual information, LP neurons projecting to AL are informative about both optic flow and running speed. To explicitly depict the degree to which different boutons integrate visual and motor signals, we computed a selectivity index, where values of −1 or 1 denote boutons modulated by only one variable (running or optic flow speed, respectively), while values close to 0 denote that the activity of a bouton was equally well correlated with both variables (Figure 6K, see Methods). As expected, selectivity indices of LP boutons in AL were distributed around 0 (median 0.18), significantly different from the distribution of V1 boutons in AL, which was biased towards 1 (median 0.58, p < 10^−27^). Therefore, V1 and LP convey different information to higher visual area AL in behaving animals: while V1 projections carry predominantly optic flow signals, the transthalamic pathway integrates these visual motion signals with information about the animals’ own movement speed. In addition, we performed similar analyses on neurons in cortical area AL recorded during the same experimental conditions. The activity of most AL neurons was modulated by both optic flow and running speed, similar to their inputs from LP, but not from V1 (Figure 6H-K). Both AL neurons and LP boutons in AL showed positive correlations with optic flow speed, but negative correlations with running speed, implying that they are activated by visual flow but suppressed by locomotion. These response characteristics could give rise to the suppression of running-induced optic flow, suggesting that visual area AL and LP-to-AL thalamocortical circuits may be specialized to process visual motion relative to self-motion.

## Discussion

We studied the anatomical and functional organization of higher-order thalamic circuits in the visual system. We found that LP neurons are part of feedforward transthalamic pathways that integrates signals from V1 with input from a large number of cortical and subcortical regions. These pathways convey target-specific visuo-motor information to different higher visual areas, distinct from the visual signals carried by V1 projections to the same cortical target, highlighting the functional difference between transthalamic and intracortical pathways. LP input to higher visual areas combines information about the visual input with the motor context it is encountered in. Transthalamic pathways may therefore link sensory information with behavioral context.

### LP is a key node of feedforward transthalamic pathways

LP neurons projecting to cortical areas AL or PM formed largely segregated populations. Yet they received relatively similar distributions of inputs from the same brain areas according to mono-synaptic rabies tracing. Both populations of LP projection neurons received input from all higher visual cortical areas, in particular from layer 6 cells. Axons from layer 6 cells form dense, small, facilitating synapses contacting distal dendrites in the thalamus and are thought to have a modulatory effect on thalamic neurons (Rockland, 1996; Alitto and Usrey, 2003; Li et al., 2003; Sherman and Guillery, 2011; Bickford, 2016; Sherman, 2016). In contrast, cortical layer 5 cells form few, but large synapses on proximal dendrites of higher-order thalamic neurons (Mathers, 1972; Li et al., 2003; Groh et al., 2014; Bickford, 2016), evoking large postsynaptic currents that can strongly influence action potential firing (Reichova and Sherman, 2004; Groh et al., 2008, 2014). Accordingly, layer 5 cells are likely to provide the main driving input from the cortex to higher-order thalamus. By far the largest number of presynaptic layer 5 cells of both AL and PM-projecting LP neurons were located in V1. These results indicate that transthalamic pathways through LP encompass a prominent feedforward component and that V1 may strongly influence the visual response properties of thalamocortical projections to higher visual areas. These findings support the long-standing hypothesis that sensory higher-order thalamic circuits form indirect, feedforward pathways (Sherman and Guillery, 2011; Sherman, 2016).

LP neurons projecting to a higher visual area did not receive input from a relatively larger number of presynaptic layer 5 or 6 cells within the same target area, arguing against strong reciprocal loops between LP and higher visual areas. This anatomical finding may appear at odds with the effects of silencing AL on the responses of LP neurons projecting to AL. While optogenetic inactivation of AL did not alter the response properties of LP boutons in AL, it did have a strong influence on their activity. Mono-synaptic rabies tracing provides an estimate of brain areas providing input to LP, but not of the functional influence of these inputs. On the other hand, optogenetic activation of parvalbumin-positive interneurons in AL likely suppressed activity in a slightly larger area (Li et al., 2019) including small parts of LM and V1, and also inhibited AL projections to other visual cortical areas, potentially leading to indirect effects on LP neurons. Moreover, the activation of PV interneurons could affect presynaptic terminals directly via GABAB receptors, reducing calcium-evoked fluorescence in LP boutons. However, this appears unlikely as the effect of inhibition was short lived (Figure 5D) in contrast to the long decay time of GABAB-induced effects (Pfrieger et al., 1994). Furthermore, thalamocortical inputs from the lateral geniculate nucleus to V1 were not suppressed via GABAB receptors in similar *in vivo* experiments (Lien and Scanziani, 2013; Gu and Cang, 2016). Finally, input from AL (and potentially other higher visual areas) may be crucial for elevating the membrane potential above spiking threshold in LP neurons and could thus be able to gate driving inputs from other sources. Importantly, however, irrespective of how much AL affects spiking in LP, our results indicate that AL does not determine the tuning properties of LP neurons projecting there, since their visual stimulus preferences were unaltered when removing AL input.

### Functional specificity of transthalamic pathways

LP conveys functionally distinct and specific information to cortical areas AL and PM. Therefore, even though some LP neurons have very large axonal projection fields (Nakamura et al., 2015; Clascá et al., 2016), LP thalamocortical projections do not broadcast identical, nonspecific signals across the cortex. These pathways carry specific visual information, tuned to spatial and temporal attributes of the visual input. Tuning curves of LP boutons for most visual stimulus features were comparable to those of V1 boutons and neurons in higher visual areas. This contrasts LP projections to V1, which convey less selective signals about the visual scene (Roth et al., 2016), and may therefore constitute a modulatory feedback pathway. While LP neurons projecting to different cortical target areas have distinct response properties, they integrate information from the same set of cortical and subcortical areas. Thus, they likely receive input from distinct sets of neurons within those areas, in particular in V1. Similarly, V1 neurons projecting to areas AL and PM constitute separate populations with distinct response properties (Glickfeld et al., 2013; Kim et al., 2018) and our data shows that the same is true for LP projection neurons. Nevertheless, most LP neurons likely project to multiple cortical areas (Nakamura et al., 2015; Clascá et al., 2016; Juavinett et al., 2020), resulting in combinatorial distribution of information along thalamocortical pathways. Determining the detailed projection motifs of single LP neurons with high-throughput methods, for instance using genetic barcoding and in situ sequencing (Chen et al., 2019), will be an essential future step towards a better understanding of higher-order thalamocortical processing.

### Transthalamic and intracortical pathways carry distinct information to the same target area

Our data indicate that LP neurons projecting to higher visual areas receive substantial input from layer 5 neurons in V1, forming indirect, transthalamic feedforward pathways (Sherman and Guillery, 2011; Sherman, 2016). These pathways could provide additional signals to higher visual areas, not present in the direct intracortical feedforward input. Consistent with this hypothesis, we find that transthalamic pathways convey information distinct from that carried by direct cortical projections from V1. LP and higher visual areas are innervated by separate populations of V1 cells, pyramidal tract and intratelencephalic neurons, respectively, with potentially different functional properties (Harris and Mrsic-Flogel, 2013). Importantly, LP neurons receive input from many other cortical and subcortical areas, notably from the superior colliculus. The superior colliculus provides particularly dense, driving input to the caudal part of rodent LP, which is mainly conveyed to lateral visual cortical areas (Zhou et al., 2017; Beltramo and Scanziani, 2019; Bennett et al., 2019), forming a second feedforward visual pathway from retina to cortex (Beltramo and Scanziani, 2019). Our study focused mainly on the anterior-lateral part of LP (Baldwin et al., 2017; Zhou et al., 2017; Bennett et al., 2019), which is innervated by the ipsilateral superior colliculus (Zhou et al., 2017, see also Figure 1), but receives its main input from visual cortical areas, providing an ideal substrate to integrate information from cortical and midbrain sources.

In our study, LP projections to higher visual areas showed a notable preference for visual stimuli with high temporal frequencies and high speed compared to V1 projections. A previous study suggested that responses to high velocity stimuli in higher visual areas are decreased after lesioning superior colliculus (Tohmi et al., 2014), indicating that the preference of LP neurons for high temporal frequency and speed could, at least in part, be inherited from the superior colliculus, and may in turn influence responses in higher visual areas. This visual channel for high temporal frequency information provided by the transthalamic LP pathway is particularly well matched with response properties of cortical area AL. AL is strongly activated by visual motion (Orsolic et al., 2019; de Vries et al., 2020) and may be specialized to process visual motion of higher temporal frequencies (Andermann et al., 2011; Marshel et al., 2011). It may therefore rely particularly strongly on the input from LP. In contrast, projections from V1 provide a visual channel with higher spatial resolution, which may be more relevant for cortical area PM which preferentially processes visual stimuli with high spatial frequency (Andermann et al., 2011; Marshel et al., 2011; Roth et al., 2012). Accordingly, our data suggests that higher visual areas integrate cortical and thalamic information differently depending on their function.

Recent studies showed that input from higher-order thalamus is critical for driving activity in motor cortex and postrhinal visual cortex (Guo Z. et al., 2017; Beltramo and Scanziani, 2019; Sauerbrei et al., 2020). To what degree pulvinar and LP affect activity in other higher visual areas is still debated (Minville and Casanova, 1998; Soares et al., 2004; Zhou et al., 2016; de Souza et al., 2020) and further studies with silencing of specific LP pathways are crucial to resolve this question.

### Transthalamic pathways through LP integrate visual and contextual information

LP not only conveys visual signals to the cortex, but also contextual information about the animals’ movement. While intracortical projections from V1 to AL carried mainly visual information about optic flow speed in animals traversing a virtual corridor, projections from LP to AL showed responses that integrated these visual signals with information about the animals’ running speed. Such motor information could originate from superior colliculus, or secondary motor and anterior cingulate cortex (Leinweber et al., 2017). Optic flow and running speed had opposing effects on the activity of LP projections to AL. These projections could therefore signal discrepancies between the expected optic flow based on the animal’s movement and the actual visual motion in the environment. Indeed, in an earlier study we showed that LP neurons projecting to V1 preferentially compute the degree of difference between running and optic flow speed (Roth et al., 2016). LP neurons projecting to V1 increase their responses with locomotion and are suppressed by optic flow. They are thus most active when an animal is running but the optic flow stops, similar to the response characteristics of a subset of V1 neurons (Keller et al., 2012). In contrast, neurons in AL and LP projections to AL are suppressed by locomotion and activated by optic flow, suggesting they contribute to processing of visual motion relative to self-motion. These neurons would be most active when the speed of optic flow is higher than expected based on the animal’s running speed, or when visual stimuli move in the environment while the animal is stationary. Therefore, LP-AL circuits could be specialized to distinguish external visual stimuli from self-generated visual feedback. Alternatively, these circuits may contribute to estimating the distance of visual stimuli during locomotion, in particular representing objects close to the animal which would result in faster optic flow during locomotion than objects further away.

One intriguing question is why transthalamic pathways would be necessary to process sensory and contextual signals, given that both visual and non-visual information can also reach cortical areas directly via cortical feedforward projections and feedback projections from higher, associative cortical regions. Our results indicate that sensory thalamocortical circuits are important to integrate cortical information with subcortical signals, in particular from the midbrain. They can thus contribute additional information that is relevant for a specific cortical area but not already present in neocortical circuits, both about specific aspects of a sensory stimulus as well as the behavioral context it is encountered in. Moreover, information about the behavioral relevance of a stimulus and the animal’s priorities, prominently represented in the superior colliculus (Krauzlis et al., 2013; Basso and May, 2017), could thus be combined with visual information from the cortex and regulate activity in higher visual areas accordingly. This hypothesis is in keeping with previous reports of a variety of non-sensory signals in pulvinar neurons, such as the animal’s focus of attention or its uncertainty about the stimulus content (Saalmann et al., 2012; Komura et al., 2013; Zhou et al., 2016; Halassa and Kastner, 2017).

Pulvinar circuits have been hypothesized to regulate communication between cortical areas (Saalmann et al., 2012; Halassa and Kastner, 2017; Jaramillo et al., 2019). Our data confirm that pulvinar circuits connect different cortical areas and show that these cortico-thalamo-cortical pathways concurrently receive signals from many other visual and non-visual brain regions. Furthermore, several long-range inhibitory circuits provide input to pulvinar neurons (Figure 1G,H) that may have the capacity to differentially regulate specific transthalamic pathways (Trageser and Keller, 2004; Halassa and Acsády, 2016; Crabtree, 2018) and thereby affect cortical information processing. We propose that higher-order thalamic circuits can both regulate sensory processing within cortical areas and control information transfer between areas (Halassa and Kastner, 2017), depending on the nature of sensory input, the behavioral circumstances, as well as the animal’s behavioral state.

## Acknowledgements

We thank Tom Otis, Alison Barth, Thomas Mrsic-Flogel, Andreas Keller and Riccardo Beltramo for comments on the manuscript. We thank Mean-Hwan Kim and Petr Znamenskyi for sharing data from CTB injection experiments, Petr Znamenskyi for code and help with the GP fit, and Rob Cambpell for setting up and help with serial-section two-photon microscopy. We thank Troy Margrie and Adam Tyson for sharing the cellfinder software and for help with automatic cell counting. We thank Andrew Murray and Molly Strom for providing rabies and AAV helper viruses and for help with optimization of rabies tracing experiments, and Niklas Schneider for help with the analysis of initial rabies results. We thank Michelle Li for animal husbandry. We thank Maxime Rio and Dylan Muir for developing software, and Thomas Mrsic Flogel, Petr Znamenskyi, Maxime Rio and Andreas Keller for discussions. We thank Ed Boyden (Chrimson), Kim Douglas and the Janelia GENIE Project (GCaMP6f), Edward Callaway (TVA), Alla Karpova and David Schaffer (retroAAV-Cre) for making their viral constructs publicly available. This work was supported by the Sainsbury Wellcome Centre Core Grant from the Gatsby Charitable Foundation and Wellcome (090843/F/09/Z), by an ERC Starting Grant (SBH), and a SNSF Project Grant (SBH).

## Author contributions

SH, MMR and AB designed the experiments. IG, FI, and MJ performed pilot experiments. IG performed rabies tracing experiments and AB conducted post-hoc tissue processing and analysis. AB, MR and IG conducted two-photon imaging and optogenetic experiments. AB, and MMR analyzed the data with help from IG. AB performed and analyzed electrophysiological recordings. SH, MMR and AB wrote the manuscript with help from IG.

## Methods

### Mice

All experiments were conducted in accordance with institutional animal welfare guidelines and licensed by the UK Home Office and the Swiss cantonal veterinary office. Mice used in this study were of either gender and were at least 6 weeks old at the start of the experiments. Mice were of the following genotype: C57BL/6j (Charles River, 42 mice for rabies tracing and two-photon imaging experiments); vGlut2-ires-cre (Vong et al., 2011, 2 mice for rabies tracing control experiments); PV-Cre (Hippenmeyer et al., 2005, 18 mice for optogenetic manipulation experiments). For more details, refer to the Reagents and Resources table.

### Surgical procedures and virus injections

Prior to surgery, mice were injected with dexamethasone (2–3 mg kg–1), atropine (0.05–0.1 mg kg–1) and analgesics (carprofen; 5 mg kg–1). General anesthesia was induced either with a mixture of fentanyl (0.05 mg kg–1), midazolam (5 mg kg–1) and medetomidine (0.5 mg.kg–1) or with isoflurane (1%–5%). All injections were made in the right hemisphere and were performed using glass pipettes and a pressure injection system (Picospritzer III, Parker). For experiments that necessitated injections into visual cortical areas AL or PM, a customized head holder was implanted using dental cement (Heraeus Sulzer or C&B), and the skull above the posterior cortex was carefully thinned and sealed with a thin layer of light-cured dental composite (Tetric EvoFlow, Ivoclar Vivadent). Intrinsic imaging maps of visual cortical areas (see Intrinsic signal imaging) were obtained several days later to identify AL and/or PM prior to injections.

For mono-synaptic rabies tracing from specific LP projection neurons, we injected a retrograde AAV-Cre (Tervo et al., 2016, ssAAV-retro/2-hSyn1-chI-iCre-WPRE-SV40p(A), 90 nl, 7.90×10^12^ vg/mL Viral Vector Facility Zurich) into either AL or PM through a small craniotomy. To mark the injection site, the pipette was coated with DiO. One week later, AAV1-Flex-nGFP-2A-G (G, Reardon et al., 2016) and AAV8-flex-GT (TVA, Wall et al., 2010, 30 nl, 1.9×10^13^ vg/mL and 1×10^14^ vg/mL, respectively) were stereotaxically injected into LP in the right hemisphere (−2.2 mm posterior to bregma, −1.6 mm lateral to bregma, −2.60 mm below the cortical surface to target AL-projecting LP neurons and −2.1 mm posterior to bregma, −1.5 mm lateral to bregma, −2.6 mm below the cortical surface to target PM-projecting LP neurons). Three days later, EnvA-pseudotyped G-deleted rabies virus (Reardon et al., 2016, CVS-N2c^ΔG^-mCherry, 60 nl, >1×10^8^ vg/mL) was injected at the same LP coordinates. The craniotomies were sealed with Tetric Evoflow light-cured dental composite. Ten to twelve days after the last injection, mice were perfused for histology (see Histology). Rabies tracing control experiments (Figure S2A-D) followed the same protocol, but no retroAAV-Cre was injected. For retrograde tracing data presented in Figure S3, fluorescent conjugate cholera toxin B (CTB; recombinant cholera toxin subunit B conjugated with Alexa fluorophores: 0.2% CTB-488 and CTB-594; Life Technologies) was injected into AL and PM.

For experiments involving two-photon calcium imaging, AAV1.hSyn.GCaMP6f.WPRE.SV40 (120 nl, 2×10^13^ vg/mL Penn Vector Core/Addgene; diluted 1:2 to 1:10 in saline) was injected either into V1, AL or PM guided by intrinsic imaging maps (see Intrinsic signal imaging) or into LP (60 nl) using stereotaxic coordinates ranging from −1.45 to −2.2 mm posterior to bregma, 1.4 to 1.6 mm lateral to bregma and 2.55 to 2.7 mm below the cortical surface. For the experiments involving optogenetic manipulations, AAV1.Syn.DIO.ChrimsonR.tdTomato (120 nl, 3.9×10^12^ vg/mL UCN, 1:5 dilution in saline solution) or AAV1.CAG.DIO.tdTomato (control, 120 nl, 2.6×10^13^ vg/mL Addgene, diluted 1:5 in saline) was injected into AL. A craniotomy of 4–5 mm diameter was made over the right hemisphere to include V1 and higher visual areas. The craniotomy was sealed with a glass coverslip and cyanoacrylate glue (UltraGel; Pattex). If not already in place from intrinsic signal imaging, a head holder was attached to the skull using dental cement (Heraeus Sulzer or C&B). Animals were given analgesics (buprenorphine 0.1 mg kg^−1^) at the end of surgery and repeatedly during recovery. Some animals additionally received antibiotics after the surgery (enrofloxacin 5 mg kg^−1^). Imaging started approximately 2 to 3 weeks after the virus injection.

### Intrinsic Signal Imaging

To determine the location of cortical visual areas AL and PM, mice underwent optical imaging of intrinsic signals (Schuett et al., 2002; Kalatsky and Stryker, 2003). Two to three days after the implantation of a head holder and thinning of the skull (see Surgical procedures), mice were initially sedated (chlorprothixene, 0.7 mg kg^−1^) and then lightly anesthetized with isoflurane (0.5–1% in O_2_) delivered via a nose cone. Visual cortex was illuminated with 700-nm light split from a LED source into two light guides. Imaging was performed with a tandem lens macroscope focused 500 μm below the cortical surface and a bandpass filter centered at 700 nm with 10 nm bandwidth (67905; Edmund Optics). Images were acquired with a rate of 6.25 Hz with a 12-bit CCD camera (1300QF; VDS Vosskühler), a frame grabber (PCI-1422; National Instruments) and custom software written in Labview (Texas Instruments). The visual stimulus was generated using the open-source Psychophysics Toolbox (Kleiner et al., 2007) based on Matlab (MathWorks) and consisted of a 25° large square-wave grating, (0.08 degrees per cycle) drifting at 4 Hz, presented on a gray background alternatively at two positions, centered at 10° elevation and either 60° or 90° azimuth. Frames in the second following stimulus onset were averaged across 16 to 32 grating presentations to generate intrinsic maps.

### Histology

Mice were euthanized with a dose of pentobarbital (80 mg kg^−1^) and transcardially perfused with 4% paraformaldehyde. Brains were extracted and post-fixed overnight in 4% paraformaldehyde. For animals who had undergone two-photon imaging, brains were embedded in 4% agarose (A9539, Sigma), cut in 200 micrometer sagittal slices on a vibratome (HM650V; Microm) and imaged on a slide scanner (Zeiss AxioScan). For animals used for anatomical tracing experiments, brains were embedded in 5% agarose and imaged using a custom-built serial-section two-photon microscope (Mayerich et al., 2008; Ragan et al., 2012). Coronal slices were cut at a thickness of 50 μm using a vibratome (Leica VT1000), and optical sections were acquired every 8 μm for rabies experiments and every 25 μm for CTB experiments. Scanning and image acquisition were controlled by ScanImage v5.6 (Vidrio Technologies, USA) using a custom software wrapper for defining the imaging parameters (Rob Campbell, 2020). For better identification of rabies virus starter cells expressing rabies, G and TVA (Figure S1D), a subset of slices were mounted in a hard-set mounting medium (2.5% DABCO (D27802; Sigma), 10% polyvinyl alcohol (P8136; Sigma), 5% glycerol, 25 mM Tris buffer pH 8.4) and imaged at higher resolution on a confocal microscope (Leica SP8).

### Two-photon Calcium Imaging

In vivo imaging experiments were performed as previously described (Roth et al., 2016). Mice were housed with an inverted light-dark cycle starting at least 5 days before the first imaging experiments. All experiments were performed during the dark phase. Animals were handled and accustomed to head restraint for 3 - 5 days. Mice were free to run on a 20-cm-diameter Styrofoam cylinder. Their running speed was measured using either an optical mouse (Logitech G700) or a rotary encoder (Kubler Encoder 1000 ppr). Imaging was performed using a commercial resonance scanning two-photon microscope (B-Scope; Thorlabs) and a Mai Tai DeepSee laser (SpectraPhysics) at 960 nm with a 16× water immersion objective (0.8 NA; Nikon). Images of 512 × 512 pixels with fields of view ranging from 180 × 180 μm to 100 × 100 μm were acquired at a frame rate of 15 or 30 Hz using ScanImage (Pologruto et al., 2003). Axonal bouton calcium measurements were performed in cortical layer 1 (62 ± 54 μm below the cortical surface). Somatic recordings were performed in layer 2/3 (166 ± 13 μm below the cortical surface). The laser power under the objective never exceeded 30mW. The surface blood vessel pattern above imaging sites was compared with the blood vessel pattern from intrinsic signal imaging maps to confirm that imaged neurons or boutons were located within a particular cortical area.

### Visual stimulation

During presentation of visual stimuli, the power supply of the monitor backlight was controlled using a custom-built circuit to present visual stimuli only at the resonant scanner turnaround points in between two subsequent imaging lines (when data were not acquired) (Leinweber et al., 2014).

#### Visual response characterization

Visual stimuli were generated using the open-source Psychophysics Toolbox (Kleiner et al., 2007) based on Matlab (MathWorks) and were presented full-field on one monitor at approximately 20 cm from the left eye of the mouse, covering 110° degrees of visual space. Visual stimuli consisted of sinusoidal gratings of all combinations of 5 different spatial frequencies (0.02, 0.04, 0.08, 0.16 or 0.32 cycles per degree) and 5 different temporal frequencies (0.5, 1, 2, 4, 8 cycles per second), presented at 4 orientations, drifting in 8 directions (0 to 360 degrees in 45 degrees increment). To avoid onset responses that would compromise the measure of temporal frequency preferences, gratings remained static for 1.2 seconds, before drifting for 2.15 seconds before the next static grating appeared. This set of 200 stimuli was randomized and presented 6 to 8 times.

#### Visuo-motor response characterization

A virtual environment consisting of a linear corridor with varying wall patterns as described previously (Poort et al., 2015; Roth et al., 2016, gratings and black and white circles on a gray background), was created in a game engine (Unity) and presented on two monitors (U2312HM; Dell) in front of the animal. The instantaneous running speed of the animal was used by a custom software written in Labview (National Instruments) to control the speed at which the animal moved through the virtual environment. Mice were habituated to this configuration for at least 2 - 3 days with 1 to 2 sessions per day. The length of the experimental session increased gradually from ~15 min to 1 hour. Mice were encouraged to run by giving them soy milk rewards through a spout either at random times or in particular corridor positions. For two-photon imaging, the optic flow was ‘uncoupled’ from the running speed, such that the animal’s locomotion did not control its movement through the virtual corridor. Instead, a movie of the virtual corridor with optic flow generated by the animal in a previous session was replayed to the mouse. Sessions in which mice showed signs that they had learned to anticipate the reward by slowing down before reward delivery, and sessions in which the median running speed was below 3 cm/s were excluded from analysis. These two criteria ensured that only recordings were included for further analysis in which animals were familiar with the virtual environment, showed a wide distribution of running speeds, and did not display stereotypical behavior.

### Optogenetic Manipulation

To silence neuronal activity in area AL, we used a 637-nm laser (Coherent) connected to a 200-micrometer optical fiber (CFMLC22, Thorlabs). The fiber was placed above AL, in between the objective used for two-photon imaging and the glass coverslip covering the craniotomy. To combine two-photon imaging and optogenetic stimulation, the laser for optogenetic stimulation was synchronized to the resonant scanner turnaround points (when data were not acquired) therefore minimizing artifacts from the monitor light (Attinger et al., 2017). The laser power was set to an average of 10 mW during stimulation. Visual stimulation was performed similarly as described above, but oblique grating orientations were excluded. Each stimulus was presented with and without laser activation and the 200 stimuli (5 spatial frequencies * 5 temporal frequencies * 4 directions * 2 laser conditions) were randomly interleaved and presented 6 to 8 times. Gratings were static for 1.2 s, before drifting for 2.3 s. When present, the laser was active for 2 s starting 0.5 s after the beginning of the static grating and 0.7 s before the onset of the drifting grating. To prevent the optogenetic manipulation during one stimulus from affecting activity in the following trial, a grey screen was displayed between stimuli for 500 ms.

### Electrophysiology

To estimate the effect of optogenetic manipulation on cortical activity, electrophysiological recordings were performed after two-photon calcium imaging in a subset of the PV-Cre mice injected with AAV-flex-Chrimson (Figure S5B,C). On the day of the recording, mice were anaesthetized under 1%–2% isoflurane, the glass coverslip covering the craniotomy was removed and the exposed cortical surface was covered with Kwik-Cast sealant (World Precision Instruments). Mice recovered from surgery for 1-2 h before the recording and were then head-fixed on a Styrofoam cylinder. The craniotomy was bathed in cortex buffer containing (in mM) 125 NaCl, 5 KCl, 10 Glucose monohydrate, 10 HEPES, 2 MgSO4 heptahydrate, 2 CaCl2 adjusted to pH 7.4 with NaOH. A silver wire was placed in the bath for referencing. One or two NeuroNexus silicon probes (A2×32-5mm-25-200-177-A64), labelled with DiI, were lowered to 600-1000 μm below the cortical surface using a micromanipulator (Sensapex). The craniotomy was then covered with 1.5% agarose in cortex buffer. Voltages from 64 or 128 channels were acquired through amplifier boards (RHD2132, Intan Technologies) at 30 kHz per channel, serially digitized and sent to an Open Ephys acquisition board via a SPI interface cable (Siegle et al., 2017). Photoactivation and visual stimulation were then performed as described above (Visual stimulation and Optogenetic manipulation).

### Data Analysis

#### CTB and mono-synaptic rabies tracing

Full resolution datasets (voxels of 2×2×8 μm for rabies experiments and 2×2×25 μm for CTB experiments) were rescaled to isometric voxels of 10 μm^3^ and registered to the Allen Mouse Common Coordinate Framework version 3 (Lein et al., 2007; Wang et al., 2020), using Elastix (Klein et al., 2010; Shamonin et al., 2014). CTB positive cells were manually counted using the cell counter plugin of Fiji (Schindelin et al., 2012; Rueden et al., 2017). For analysis of rabies virus tracing experiments, only brains were included in which we could locate starter cells within LP borders, in which G positive cells were found exclusively in LP and in which the retroAAV injection (labelled with DiO and targeted using intrinsic signal imaging) was located in AL or PM as defined by the Allen Mouse Common Coordinate Framework (Lein et al., 2007).

Fluorescent, rabies-positive cells were automatically detected using cellfinder (Adam Tyson et al., 2020) (commit 9ccc641a). Cell candidates were detected as threshold crossings on filtered images and classified as cell or non-cell by a deep neural network. The deep neural network was trained using a large dataset of manually identified cells and non-cells. Running the same automatic cell detection on control brains from Figure S2A-D yielded a low number of false-positive cells (19 and 53 cells, i.e. 1.1 +/− 1.2 % of the total number of cells detected in experimental brains), mostly corresponding to bright particles at the surface of the brain. The location of detected cells was analyzed using custom scripts in Python and figures were generated using matplotlib (Caswell et al., 2019). In the cortex, a fraction of cells was detected in the white matter just below layer 6b (see examples in Figure 1B,C and Figure S1E-G). To account for these, any cell detected in the white matter less than 50 μm from the cortical border assigned by the common coordinate framework was allocated to layer 6b of the closest cortical area. The total number of cells per brain varied from animal to animal (see Figure S2D), therefore, cell numbers per brain region are reported as proportion of detected cells per brain. Dorsal views of cortical layers (Figure 2C) are the maximum projection of each layer of interest along the dorso-ventral axis.

Dorsal projections of cell density histograms in different layers across cortical areas (Figure 2D, Figure S2H) were computed in 3D bins of 20 μm × 20 μm (along the antero-posterior and medio-lateral axis) × the thickness of the layer. Values were normalized by the total number of cells per brain and then averaged across brains. To take into account the variable layer thickness at different cortical positions, particularly in the dorsal part of the brain where the layers are tangential to the dorso-ventral axis, each bin was divided by its volume. Rabies-virus positive cell density is therefore expressed as percentage of total cells per cubic millimeter. Transverse views of cell density histograms (Figure S3D-F) were computed in 3D bins of 10 μm × 10 μm × 200 μm and upsampled to 5 μm² pixels with linear interpolation using scikit resize (Walt et al., 2014).

#### Two-photon imaging

Image stacks were processed using custom-written scripts in Matlab (Mathworks) as described in Roth et al 2016. Briefly, to correct for x-y motion, two-photon imaging frames were registered to a 30-frame average using a phase-correlation algorithm. Frames with large motion were detected by inspecting the registration displacement results and were subsequently discarded from further analysis. Regions of interest (ROIs) were detected semi-automatically using intensity thresholding combined with PCA-ICA refinement and validated and refined manually. All time-series were extracted and analyzed with custom written functions using the TimeSeriesAnalysis package (Muir et al., 2020) (see Table) For recordings of neuronal somata, contaminating signals coming from densely labelled neuropil were subtracted using an Asymmetric Student-t model (ast_model available here: https://github.com/BaselLaserMouse/ast_model). ΔF/F calcium transients were obtained by using the 25th percentile over the entire fluorescence distribution as F0. Firing rates per imaging frame were then inferred from ΔF/F using a compressive sensing technique (Dyer et al., 2013; Roth et al., 2016).

#### Electrophysiology

Spikes were sorted with Kilosort (Pachitariu et al., 2016) using procedures previously described (Chabrol et al., 2019). Each unit was attributed to the channel on which the extracellular waveform had the highest amplitude. Recording depth was estimated based on the DiI track and only channels in close proximity (< 100 um) to the Chrimson injection site were included for analysis. Single units with average firing rate significantly higher in laser trials than in controls trials (putative PV+ interneurons expressing chrimson) were excluded from the analysis.

#### Analysis of visual responses

The response to each stimulus was measured as the inferred firing rate averaged over a window starting 250 ms after the onset of grating movement and ending either at the end of the stimulus presentation (Figure 3 and 4) or at the end of the laser stimulation (Figure 5). Responses were then fitted using a gaussian process (GP) regression model as previously described (Kim et al., 2018). The GP fit has several advantages compared to more classical parametric methods: (1) it does not assume independence between the different stimulus dimensions (e.g. between spatial and temporal frequency tuning), (2) it does not constrain the shape of the response profile (for instance to be Gaussian) but only assumes that response variations are continuous; (3) it is probabilistic and therefore provides not only an estimate of the average response but also of its variance; (4) it easily allows for integration of other parameters that influence neuronal activity but are harder to include in parametric fits, such as the running speed of the animal. GP predictions were made from five predictors: the spatial frequency, the temporal frequency and the direction of the stimulus, the average running speed of the animal during stimulus presentation and the presence of the laser (0 for control trials and 1 for laser trials).

As previously described (Kim et al., 2018), the GP regression model is fitted by estimating the parameters of the kernel function (x_i_, x_j_), which defines the covariance of the neuronal activity as a function of the similarity of stimuli x_i_ and x_j_, defined by their respective spatial frequencies SF, temporal frequencies TF, directions θ, running speed s and the presence of laser L. We used a product of a squared exponential (SE) kernel for spatial and temporal frequencies and a periodic kernel for direction:

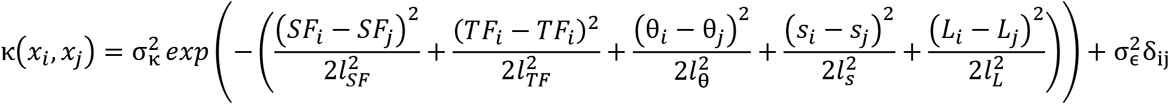

Length scale parameters l_SF_, l_TF_, l_θ_, l_s_, and l_L_ determine how quickly (x_i_, x_j_) declines with changes of that stimulus dimension. Variance parameters 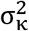 and 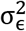 correspond to the stimulus-dependent and stimulus-independent (i.e., noise) components of the response. δij is Kronecker delta and is one if i = j and zero otherwise. Optimization is accomplished by maximizing the likelihood p(r|X) of observed responses r given the set of stimuli X.

The GP model was implemented in Python using the gpflow library (Matthews et al., 2017). We used Gamma(2,1) as a prior for length scale parameters l_SF_, l_TF_, l_θ_, l_s_, and l_L_. In addition, to avoid overfitting, we constrained l_SF_, l_TF_, l_s_, and l_L_ ≥ 0.25. After optimizing the kernel parameters, we searched for the stimulus that evoked the maximum response using the Nelder-Mead method of the scipy minimize function. We then defined a signal to noise ratio (SNR) as:

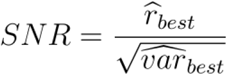

where r_best_ and var_best_ are the predicted mean and variance of the response to the best stimulus. ROIs were considered responsive if the signal to noise ratio was above two and the R^2^ of the fit, defined as R^2^ = 1 − Σ_i_((r_i_−p_i_)^2^)/Σ_i_((r_i_ − r)^2^) where r_i_ is the response at stimulus *i* and p_i_ is the prediction of the GP fit for the same stimulus, was above 0.1. All results presented in the paper could be qualitatively reproduced using a parametric fit instead of the GP fit.

Determining a preferred stimulus feature of a neuron (e.g. preferred spatial frequency) is only meaningful if its response is significantly modulated across this stimulus dimension. When reporting preferred visual stimuli (Figures 3,4,5,S4,S5), we therefore only included ROIs for which the predicted response to the preferred stimulus, r_best_, was at least 1.33 standard deviations above that to the stimulus evoking the smallest response along that dimension r_worst_ (e.g. for responses to different spatial frequencies for stimuli with the same temporal frequency and grating direction,):

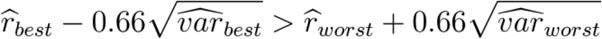

To estimate the preferred stimulus, the search of the maximum of the GP fit was bounded to the range of presented spatial and temporal frequencies ([0.02 - 0.32] and [0.5 - 8]). For Figure 5 and Figure S5 the same search was performed with the laser parameter L fixed to 0 for estimating the preferred stimulus in control trials and fixed to 1 for laser trials. Preferred speed was defined as the ratio between the preferred temporal and spatial frequency. To average across boutons/neurons or display them on the same color scale, responses were z-scored using the mean and standard deviation of the inferred spike rate. Response matrices (Figures 3–5) were obtained by averaging the z-scored response amplitude of all visually responsive ROIs for every combination of spatial and temporal frequencies at the preferred direction of each ROI. The similarity between such matrices was then evaluated by computing the Pearson correlation coefficient between the average matrix of each individual imaging session and the average matrix of all AL neurons (Figure 3G), of all PM neurons (Figure 4D) or of responses in control trials of all LP boutons in AL that were inhibited by laser stimulation (Figure 5H).

To measure response specificity to different visual stimulus features (Figure S4I-K), we computed tuning curves of predicted responses to varying spatial or temporal frequencies while keeping all other stimulus parameters fixed to those evoking the peak response. We then measured the full-width half maximum of the resulting curve (examples of such curves for individual neurons can be found in Figure 3C).

To quantify the effect of optogenetic manipulation on LP boutons, we included all visual stimuli that evoked a response with SNR of the GP prediction above 2, and with an amplitude of at least ⅔ of the response to the best stimulus. Responses of individual trials to these stimuli were pooled to test the effect of laser stimulation using a Wilcoxon rank-sum test (Trials with and without laser are not paired). Boutons were defined as significantly suppressed if their average response was significantly lower in laser trials than in control trials (alpha < 0.05). To compare tuning curves in response to different visual stimulus properties with and without optogenetic laser stimulation, we included boutons that were significantly responsive in both conditions, defined as SNR of the response to the preferred stimulus above 2. Tuning curves of individual boutons were plotted centered on and relative to the preferred stimulus (Figure S5J) and included responses at the preferred grating direction and the preferred temporal or spatial frequency for spatial and temporal frequency tuning curves, respectively.

#### Analysis of visuo-motor responses

To identify responsive boutons or neurons, we measured the skewness of ΔF/F values of individual ROIs over the recording. ROIs with skewness > 1 were considered to be responsive. For each responsive bouton or neuron, a normalized cross-correlation was computed by obtaining time-dependent Pearson correlation coefficients between its inferred spike rate and a behavioral variable (running speed or optic flow speed resampled at the imaging frame rate) over a range of lags between −1 to 1 second (corresponding to 60 different lags with 30hz imaging frame rate). For each behavioral variable and each bouton or neuron, we then determined the lag with the highest absolute correlation coefficient. From these values, we computed the median lag of the neuronal population as m_Lag_ (separately for running speed and optic flow speed; Figure S6). We computed a mean cross-correlation coefficient for each bouton or neuron and each behavioral variable (R_RS_ for running speed and R_OF_ for optic flow speed) by averaging the time-dependent Pearson coefficients over lags in a window of 250 ms centered on m_Lag_. To determine if a bouton or a soma was correlated with a behavioral variable, we defined a circular threshold: the magnitude of the vector |R| composed by [R_OF_, R_RS_] was computed as the square root of the sum of squared R_OF_ and squared R_RS_. Only boutons with |R| >= 0.1 were included in the following analysis. The interaction angle θ was determined using the mean cross-correlation coefficients, and was computed as θ = atan(R_RS_/R_OF_). For estimating the circular median interaction angle per session, we computed θ_pop_ = atan(R_RSpop_/R_OFpop_) where R_RSpop_ and R_OFpop_ are the median R_RS_ and R_OF_ across boutons or somata. For each bouton or soma, we calculated a selectivity index as the difference between absolute R_OF_ and absolute R_RS_ divided by the sum between absolute R_OF_ and absolute R_RS_.

### Statistics

We used two-sided Wilcoxon rank-sum tests for independent group comparisons, and two-sided Wilcoxon signed-rank tests for paired tests. We used circular statistics and circular metrics (Berens, 2009) when required (Figure 6J). Test were performed using either matlab or rpy2. No statistical methods were used to pre-determine experimental sample sizes.

### Acronyms

| ACRONYM | FULL NAME | ACRONYM | FULL NAME |
| --- | --- | --- | --- |
| a.u. | Arbitrary units | PM | Posteromedial visual area |
| AAV | Adeno-associated virus | POL | Posterior limitans nucleus of the thalamus |
| ACC | Anterior cingulate cortex | POR | Postrhinal area |
| AL | Anterolateral visual area | PPT | Posterior pretectal area |
| AM | Anteromedial area | PV | Parvalbumin |
| APN | Anterior pretectal nucleus | RHP | Retrohippocampal region |
| Apo | Posterior auditory area | RL | Rostrolateral visual area |
| AUD | Auditory areas | RN | Red nucleus |
| CTB | Cholera toxin B | RS | Running speed |
| CTX | Cortex | RSP | Retrosplenial cortex |
| DAPI | 4',6-diamidino-2-phenylindole | RSPagl | Agranular retrosplenial cortex |
| dLGN | Dorsal lateral geniculate nucleus | RSPd | Dorsal part of the retrosplenial cortex |
| ECT | Ectorhinal areas | RSPv | Ventral part of the retrosplenial cortex. |
| FWHM | Full width half maximum | RT | Reticular thalamic nucleus. |
| G | Rabies virus glycoprotein | SCdg | Superior colliculus, deep gray layer |
| GP | Gaussian Process | SCdw | Superior colliculus, deep white layer |
| HVA | Higher visual areas | SCig | Superior colliculus, intermediate gray layer |
| IGL | Intrageniculate leaflet | SCiw | Superior colliculus, intermediate white layer |
| INC | Interstitial nucleus of Cajal | SCop | Superior colliculus, optic layer |
| LD | lateral dorsal nucleus of the thalamus | SCsg | Superior colliculus, superficial gray layer |
| LI | Laterointermediate area | SCzo | Superior colliculus, zonal layer |
| LM | Lateromedial area | SF | Spatial frequency |
| LP | Lateral posterior nucleus of the thalamus | SS | Somatosensory areas |
| M1 | Primary motor cortex | SubG | Subgeniulate nucleus |
| M2 | Secondary motor cortex | TEa | Temporal association area |
| MPT | Medial pretectal area | TF | Temporal frequency |
| MRN | Midbrain reticular nucleus | TVA | Tumor virus A protein |
| ND | Nucleus of Darkschewitsch | V1 | Primary visual cortex |
| NOT | Nucleus of the optic tract | VGLUT2 | Vesicular glutamate transporter 2 |
| NPC | Nucleus of the posterior commissure | vLGN | Ventral lateral geniculate nucleus |
| OF | Optic Flow | WT | Wild type |
| OP | Olivary pretectal nucleus | ZI | Zona incerta |
| PL | Prelimbic area |  |  |

### Reagents and Resources Table

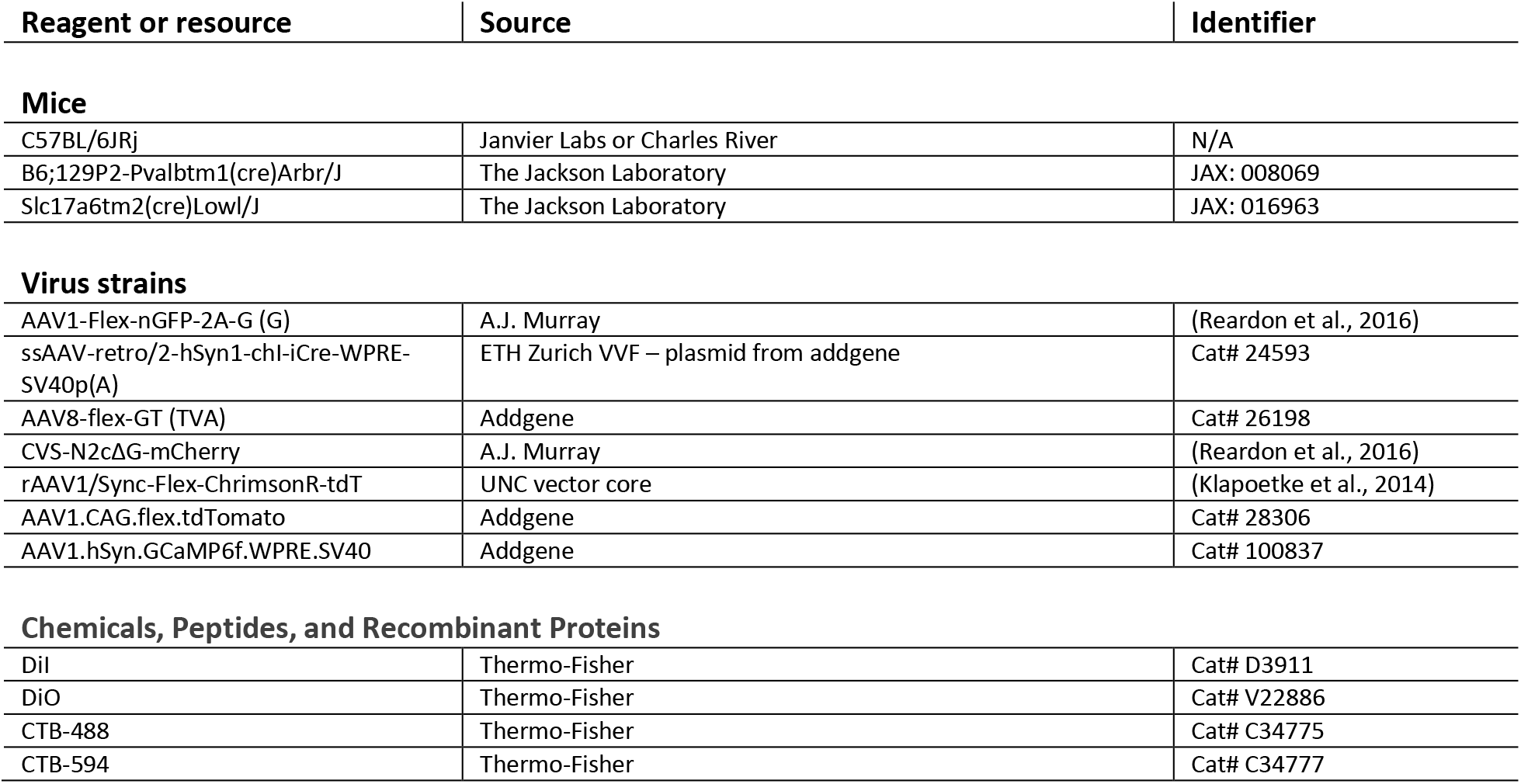
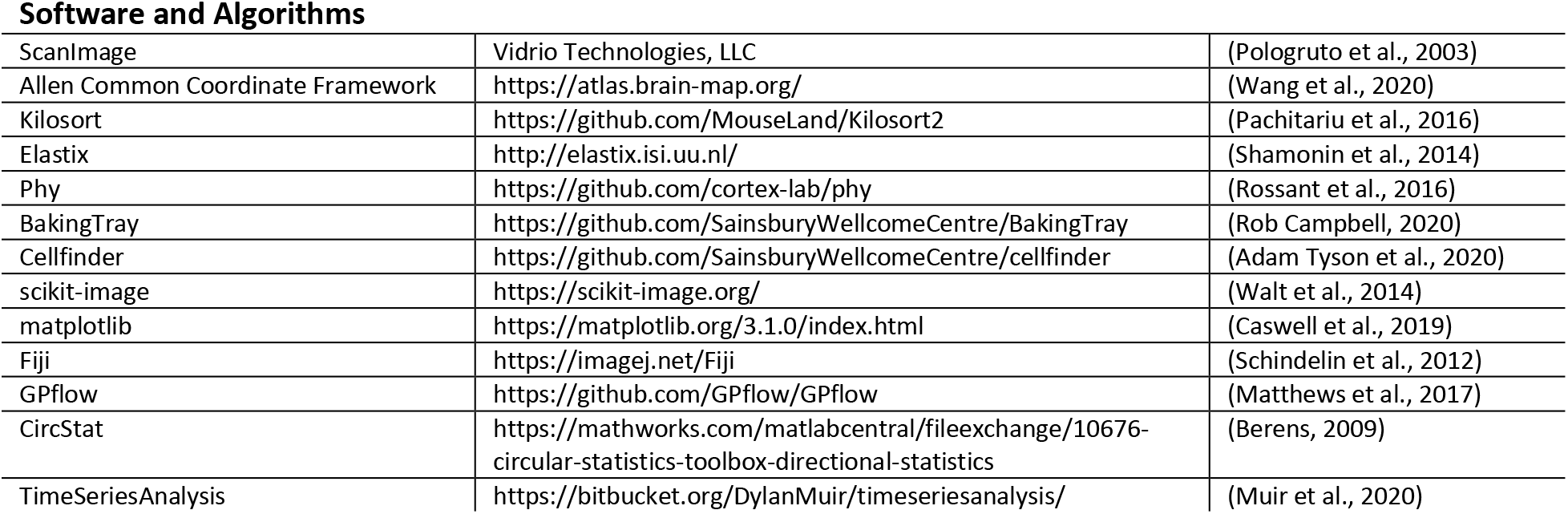

---

**Figure S1.**
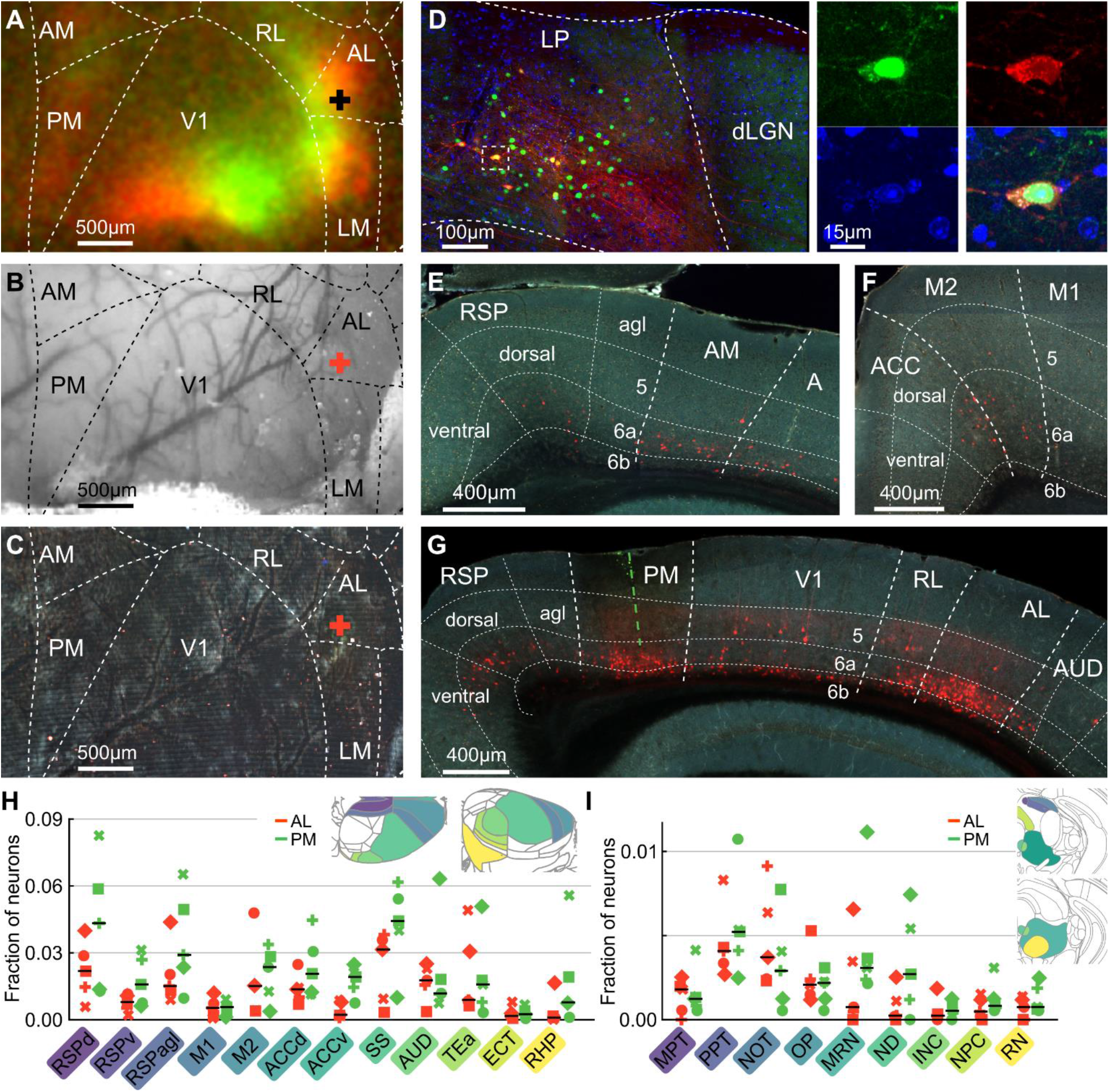
**(A)** Example intrinsic signal imaging map used to identify and target higher visual area AL for the retroAAV-Cre injection. Intrinsic responses evoked by two spatially separated visual stimuli (see Methods) are color-coded in green and red. The cross marks the injection site. AL: anterolateral area, AM: anteromedial area, LM: lateromedial area, PM: posteromedial area, RL: rostro lateral area, V1: primary visual cortex. **(B)** Surface blood vessel pattern corresponding to the intrinsic imaging map in (A). **(C)** Dorsal view of the same area shown in (A) and (B) in the perfused brain reconstructed after serial-section two-photon imaging. The brain was registered to the Allen common coordinate framework and area borders were aligned to the intrinsic map in (A) using the blood vessel pattern in (B). **(D)** Left: confocal image of a coronal slice through the lateral posterior thalamic nucleus (LP) injection site, showing starter neurons expressing G, TVA and rabies virus (green and red overlap), and neurons expressing G and TVA only (green). Dashed lines indicate the borders of LP and the dorsal lateral geniculate nucleus (dLGN). Right: magnified image of a rabies starter cell in LP. Nuclear green label indicates the presence of G protein, cytosolic green label reflects TVA expression and red label indicates the presence of rabies virus. DAPI staining is shown in blue. **(E,F)** Example images of coronal slices showing rabies-labelled cells (red) presynaptic of AL-projecting LP neurons after retroAAV injection into AL. A: anterior visual area, ACC: anterior cingulate cortex, agl: agranular, AM: anteromedial area, M1: primary motor cortex, M2: secondary motor cortex, RSP: retrosplenial cortex. **(G)** Example image of a coronal slice showing the cortical injection site of retroAAV-Cre in PM (pipette track marked in green) and rabies-expressing cells (red) presynaptic to PM-projecting LP neurons. AL: anterolateral area, agl: agranular, AUD: auditory areas, PM: postromedial area, RL: rostrolateral area, RSP: retrosplenial cortex, V1: primary visual cortex. **(H)** Fraction of rabies-positive cells presynaptic to AL- (orange) and PM-projecting (green) LP neurons across cortical areas not detailed in Figure 1. ACCd: anterior cingulate cortex, dorsal part, ACCv: anterior cingulate cortex, ventral part, AUD: auditory areas, ECT: ectorhinal areas, M1: primary motor cortex, M2: secondary motor cortex, RHP: retrohippocampal region, RSPagl: agranular part of the retrosplenial cortex, RSPd: dorsal part of the retrosplenial cortex, RSPv: ventral part of the retrosplenial cortex, SS: somatosensory areas, TEa: temporal association areas. **(I)** Fraction of rabies-positive presynaptic cells across midbrain areas, excluding the anterior pretectal nucleus and the superior colliculus (presented in Figure 1). INC: interstitial nucleus of Cajal, MPT: medial pretectal area, MRN: midbrain reticular nucleus, ND: nucleus of Darkschewitsch, NOT: nucleus of the optic tract, NPC: nucleus of the posterior commissure, OP: olivary pretectal nucleus, PPT: posterior pretectal area, RN; red nucleus.

**Figure S2.**
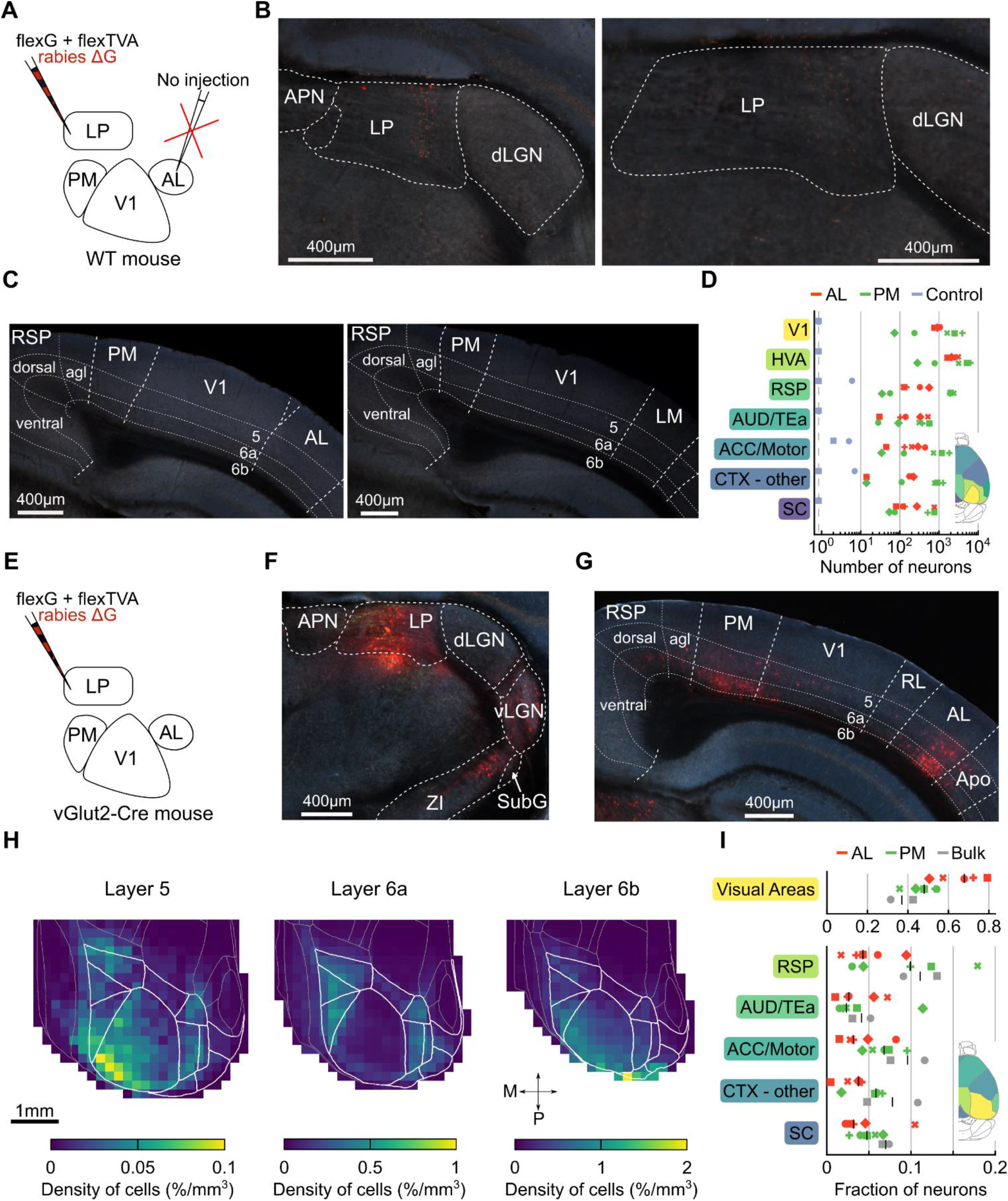
**(A)** Design of rabies control experiment. The experimental protocol was similar to the one described in Figure 1, but no retroAAV-cre was injected. WT: wild-type **(B)** Coronal sections through LP of two different animals showing no G or TVA expression. **(C)** Coronal sections through the cortex from the same animals presented in (B) showing the absence of rabies-positive presynaptic cells. **(D)** Numbers of rabies-positive cells detected by the automated cell counting software across brain areas in brains injected with retroAAV in AL (orange) in PM (green), and without injection of retroAAV-Cre (blue). Markers represent single animals. Dashed line indicates 0. **(E)** Schematic of the experimental design. To label cells presynaptic to LP neurons without specific projection target, we injected AAV-flex-G, AAV-flex-TVA and rabies virus into LP of VGluT2-Cre mice. **(F)** Coronal slice through the LP injection site showing rabies-positive cells in subcortical areas, presynaptic to LP neurons without specific projection target. **(G)** Coronal slice showing rabies-positive cells presynaptic to LP neurons in cortical areas. **(H)** Dorsal view of the average relative density of cells presynaptic to LP neurons per volume (see Methods) in layer 5 (left), layer 6a (middle) and layer 6b (right); 2 mice. White lines indicate the border of cortical areas as in Figure 2A. **(I)** Fraction of rabies-positive cells presynaptic to AL-projecting LP neurons (orange), PM-projecting LP neurons (green), and the general LP population (grey). Markers represent single animals. Bottom right corner: dorsal view of color-coded cortical areas. ACC/Motor: anterior cingulate areas and motor areas, agl: agranular, AL: anterolateral area, APN: anterior pretectal nucleus, Apo: posterior auditory area, AUD/TEa: auditory areas and temporal association areas, CTX: cortex, dLGN: dorsal lateral geniculate nucleus, HVA: higher visual areas, LM: lateromedial area, LP: lateral posterior nucleus, PM: posteromedial area, RL: rostrolateral area, RSP: retrosplenial cortex, SC: superior colliculus, SubG: subgeniculate nucleus, V1: primary visual area, vLGN: ventral lateral geniculate nucleus, WT: wild type. ZI: zona incerta.

**Figure S3.**
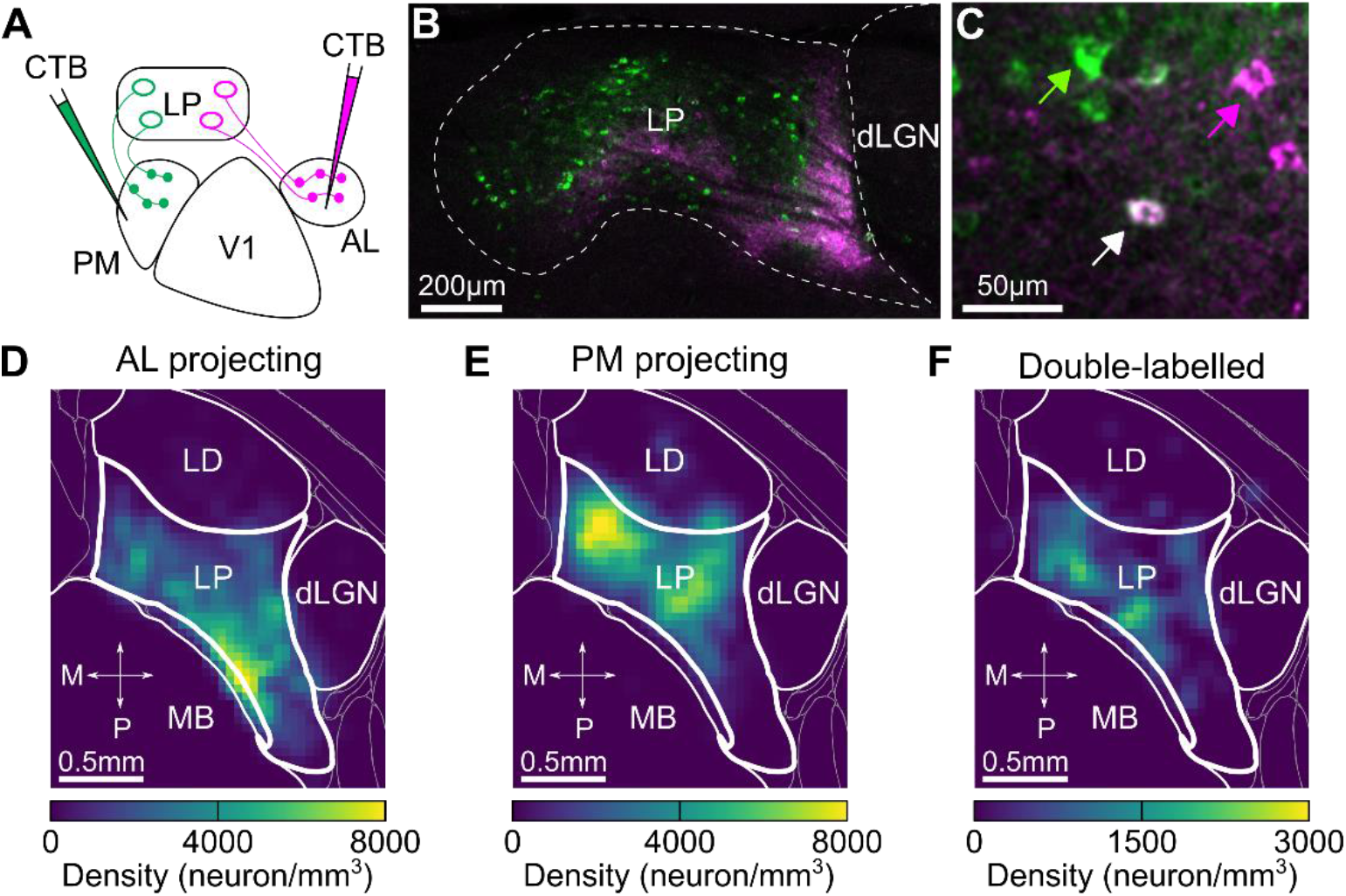
**(A)** Schematic of the experimental design. To determine the extent of overlap between AL and PM-projecting LP neurons, retrograde tracers of different colors (CTB) were injected into AL and PM. **(B)** Coronal section through LP showing the spatial distribution of PM- (green) and AL-projecting (pink) neurons. **(C)** Magnified area from (B). Arrows indicate examples of a PM-projecting LP neuron (green), an AL-projecting neuron (pink), and a neuron that targets both PM and AL (double-labelled, white). **(D-F)** Dorsal view of the average relative density of AL-projecting (D, 3504 and 2102 neurons), PM-projecting (E, 2123 and 2406 neurons), and double-labelled neurons in LP (F, 442 and 412 neurons) neurons in LP. Data from two mice. AL: anterolateral area, CTB: choleratoxin subunit B, dLGN: dorsal lateral geniculate nucleus, LD: lateral dorsal nucleus, LP: lateral posterior nucleus, M: medial, P: posterior, V1: primary visual cortex.

**Figure S4.**
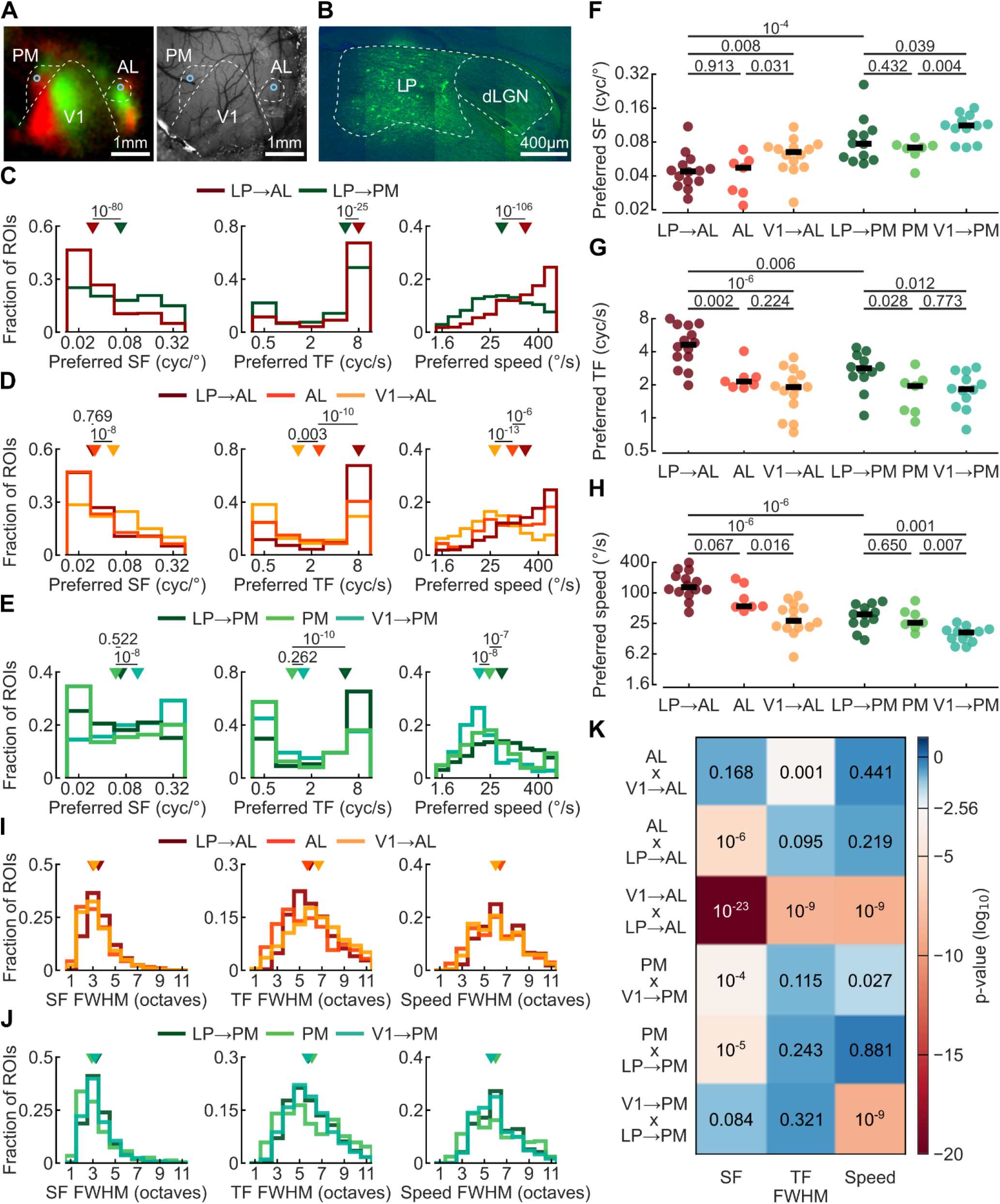
**(A)** Example intrinsic imaging map (left) showing responses to two spatially separated visual stimuli in green and red, and image of blood vessel pattern of the imaging area under the cranial window (right). Blue circles indicate the two imaging sites recorded in the posteromedial area (PM) and the anterolateral area (AL). **(B)** Example image of GCamP6f injection site. dLGN: dorsal lateral geniculate nucleus, LP: lateral posterior nucleus. **(C)** Distribution of preferred spatial frequency (left, 2237 LP boutons in AL and 2059 LP boutons in PM), preferred temporal frequency (middle, 1333 and 1128 LP boutons for AL and PM) and preferred speed (right, 2468 and 2342 boutons for AL and PM respectively) of grating stimuli for significantly modulated LP boutons recorded in AL (dark red) and in PM (green). Triangles indicate the median. **(D)** Same as (C) for AL neurons (orange, 172, 126, and 231 neurons for SF, TF and speed respectively), LP boutons (dark red) and V1 boutons (yellow, 2555, 1637, and 2928 boutons for SF, TF and speed respectively) recorded in AL. **(E)** Same as (C) for PM neurons (green, 214, 181, and 288 neurons for SF, TF and speed respectively), LP boutons (dark green) and V1 boutons (blue, 2327, 1659, and 2535 boutons for SF, TF and speed respectively) recorded in PM. **(F)** Median preferred spatial frequencies of grating stimuli per imaging session of LP boutons in AL and PM, AL and PM neurons, and V1 boutons in AL and PM. Circles represent individual sessions. Black dashes indicate the median of all sessions. **(G)** Same as (F) for preferred temporal frequencies of grating stimuli. **(H)** Same as (F) for preferred grating speed. **(I)** Distribution of full width half maxima (FWHM) of GP fit prediction (see Methods) of spatial frequency (SF) tuning curves at the preferred temporal frequency and preferred direction of the grating (left), of temporal frequency (TF) tuning curves at the preferred spatial frequency and preferred direction (middle), and of speed tuning curves at the preferred grating direction (right). **(J)** Same as (I) for full width half maxima of tuning curves of LP boutons in PM, PM neurons and V1 boutons in PM. **(K)** P-values of Wilcoxon rank-sum test for relevant pairwise comparisons of tuning width, color coded by significance level (threshold adjusted for multiple comparison using Bonferroni correction). In all panels data from 7 sessions in 5 mice for AL neurons, 8 sessions in 5 mice for PM neurons, 14 sessions in 14 mice for LP boutons in AL, 12 sessions in 10 mice for LP boutons in PM, 14 sessions in 7 mice for V1 boutons in AL and 12 sessions in 7 mice for V1 boutons in PM.

**Figure S5.**
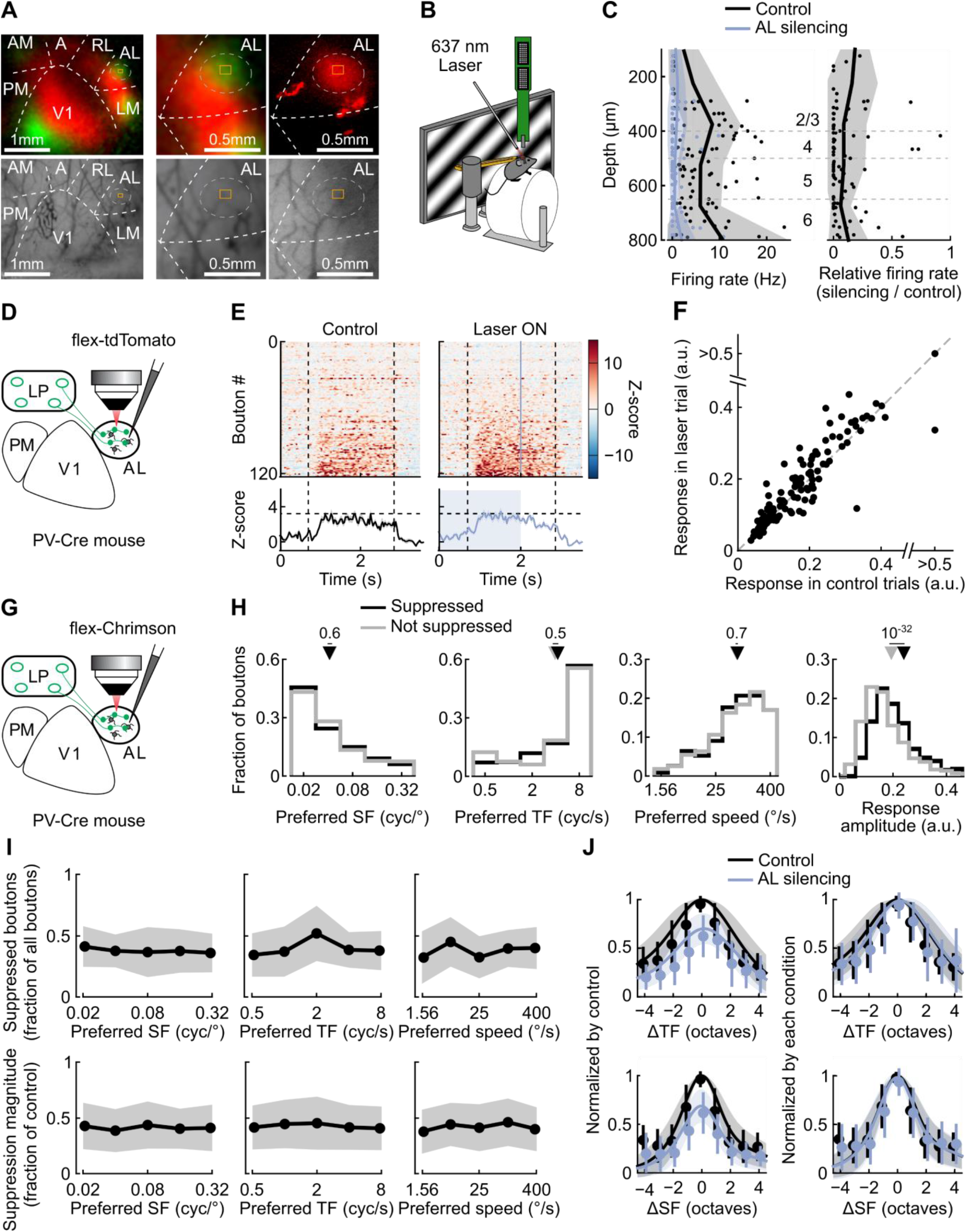
**(A)** Example intrinsic imaging map obtained to determine the location of cortical area AL for two-photon imaging and injection of AAV-flex-Chrimson. Top left: intrinsic imaging map showing responses to two spatially separated visual stimuli (red and green). Middle: Zoomed-in cutout from image on the left. Top right: image showing the extent of Chrimson expression (red) in AL, approximated by the dotted circle. Red specks outside the circle are imaging artifacts. Bottom: surface blood vessel pattern of the imaging sites depicted on top. Yellow square indicates the imaging site. **(B)** Schematic of the experimental design for electrophysiological recordings. The cranial window was removed, and a multi-channel silicon probe was inserted into the cortical site with Chrimson expression. Awake mice were presented with gratings and a 637-nm laser was used to activate Chrimson-expressing PV neurons to silence cortical activity similar to experiments in Figure 5. **(C)** Mean firing rate during visual stimulation with (blue) and without (black) laser (left) and proportion of remaining response in laser trials (right) as a function of recording depth for single units not excited by the laser. Dots depict individual units. Shading indicates standard deviation. 42 single units, 10 mice. **(D)** Schematic of the laser control experiment. PV-Cre mice were injected with AAV-flex-tdTomato in AL and GCaMP6f in LP. Visually-evoked activity of LP boutons in AL was imaged with and without laser stimulation over AL. **(E)** Top: time course of z-scored neuronal activity of individual boutons. For each bouton, activity was averaged across grating stimuli evoking a response (see Methods) in control trials (left) and laser trials (right). Responses are aligned to the onset of the laser. Blue shading indicates time of laser stimulation. Dashed lines show duration of moving grating. Bottom: Average response across all boutons. Grey shading indicates sem. 123 boutons from 4 sessions in 4 mice **(F)** Relationship between the average response of individual boutons with and without laser stimulation. **(G)** Schematic of the experimental design, as in Figure 5A. **(H)** Distribution of preferred stimulus properties of suppressed (black, 610, 337, 731, and 973 boutons for SF, TF, speed and amplitude respectively) and non-suppressed (grey, 694, 461, 990, and 1326 boutons for SF, TF, speed and amplitude respectively) LP boutons. Triangles indicate medians. **(I)** Fraction of significantly suppressed boutons per session (top, 13 sessions), and magnitude of suppression (bottom, 973 boutons) as a function of the preferred spatial frequency (left), temporal frequency (middle) and speed (right). Shading indicates standard deviation across sessions (top) or boutons (bottom). **(J)** Left: Mean response curves to varying temporal (top) and spatial (bottom) grating frequencies of boutons significantly, but not fully suppressed during AL silencing (see Methods, 169 and 257 boutons for TF and SF) in control trials (black) and laser trials (blue), normalized by the response to the preferred stimulus in control trials, plotted centered on and relative to the preferred frequency. Right: same as on the left, but normalized by the maximum response within each condition. Dots and error bars indicate means and the standard deviation of the raw data across boutons, while the lines and shading indicate means and standard deviation obtained from the prediction of the GP fit (see Methods).

**Figure S6.**
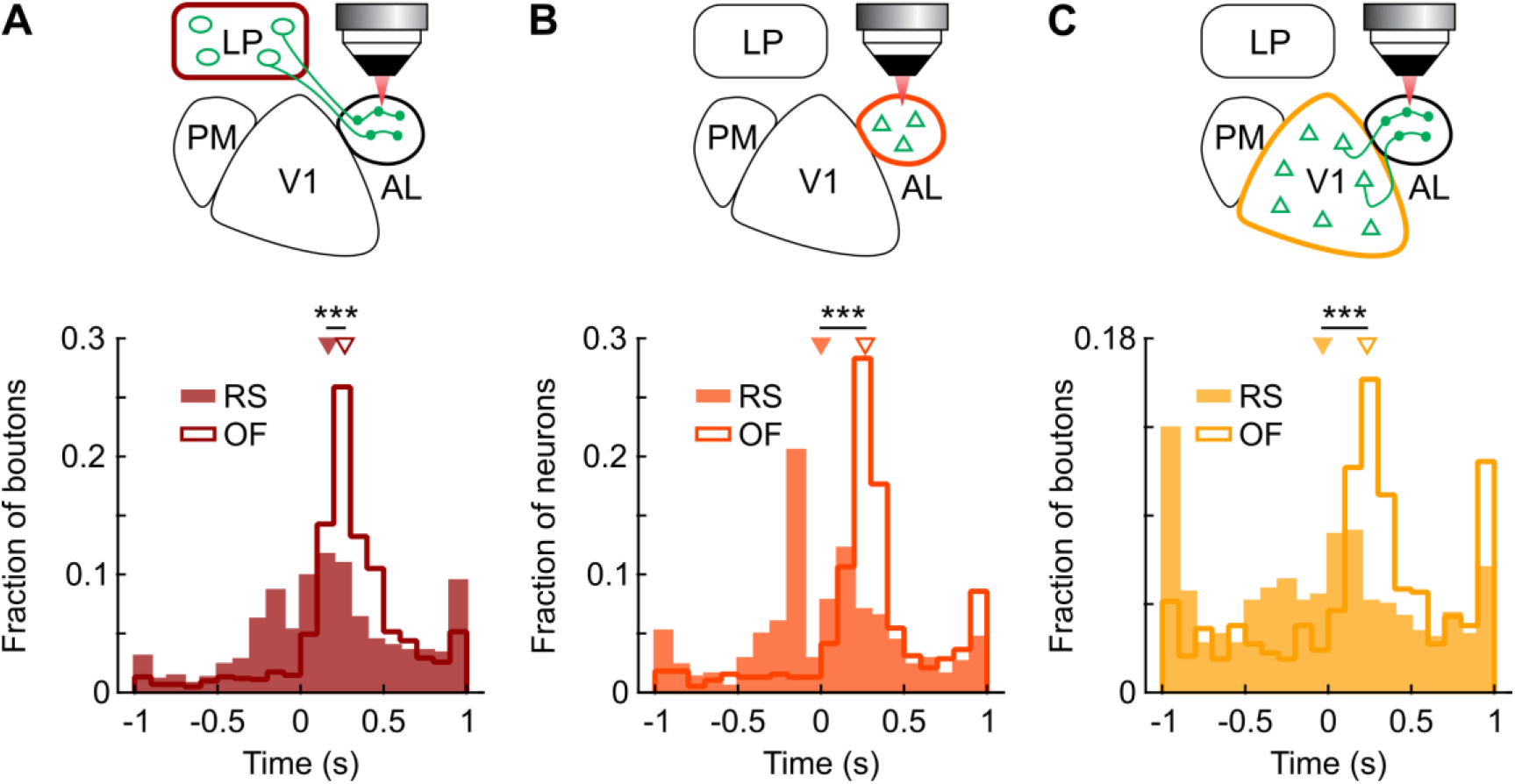
**(A,B,C)** Top: schematic of the experimental design. Bottom: distribution of lag times between running speed (RS, filled histogram) or optic flow (OF, line) and the activity of LP boutons imaged in AL (A, 1437 boutons with correlation coefficients above the circular threshold of 0.1 from 43 sessions in 18 mice), AL neurons (B, 385 neurons with correlation coefficients above the circular threshold of 0.1 from 15 sessions in 5 mice) and V1 boutons in AL (C, 647 boutons with a correlation above the circular threshold of 0.1 from 6 sessions in 3 mice) for which the absolute cross-correlation was maximal. *: P < 10^−3^; **: P < 10^−10^; ***: P < 10^−15^.

